# Development of an ASO therapy for Angelman syndrome by targeting an evolutionarily conserved region at the start of the *UBE3A-AS* transcript

**DOI:** 10.1101/2021.07.27.453820

**Authors:** Scott V. Dindot, Sarah Christian, William J. Murphy, Allyson Berent, Jennifer Panagoulias, Annalise Schlafer, Johnathan Ballard, Kamelia Radeva, Ruth Robinson, Luke Myers, Thomas Jepp, Hillary Shaheen, Paul Hillman, Kranti Konganti, Andrew Hillhouse, Kevin R. Bredemeyer, Lauren Black, Julie Douville, the FIRE consortium

## Abstract

Angelman syndrome is a devastating neurogenetic disorder for which there is currently no effective treatment. It is caused by mutations or epimutations affecting the expression or function of the maternally inherited allele of the ubiquitin-protein ligase E3A (*UBE3A*) gene. The paternal *UBE3A* allele is imprinted in neurons of the central nervous system (CNS) by the *UBE3A* antisense (*UBE3A-AS*) transcript, which represents the distal end of the *SNHG14* transcription unit. Reactivating the expression of the paternal *UBE3A* allele in the CNS has long been pursued as a therapeutic option for Angelman syndrome. Here, we designed and optimized antisense oligonucleotides (ASO) targeting an evolutionarily conserved region demarcating the start of the human *UBE3A-AS* transcript and show that ASOs targeting this region can reverse imprinting of *UBE3A* in cultured Angelman syndrome neurons and throughout the CNS of a non-human primate model. Findings from this study advanced the first investigational molecular therapy for Angelman syndrome into clinical development (ClinicalTrials.gov, NCT04259281).

**SUMMARY:** Here, we describe the preclinical studies supporting the first investigational molecular therapy for Angelman syndrome to advance into clinical development (ClinicalTrials.gov, NCT04259281).

## INTRODUCTION

Angelman syndrome is a devastating neurogenetic disorder caused by mutations or epimutations leading to the loss of function or expression of the maternal allele of the ubiquitin-protein ligase E3A (*UBE3A*) gene [1, 2]. The prevalence of Angelman syndrome is estimated at approximately 1 in 12,000 to 20,000 live births, with males and females affected similarly [3]. Individuals with Angelman syndrome have developmental delay, severe motor and cognitive deficits, epilepsy, and a range of other neurological symptoms [4, 5]. They have an average life span and require life-long, full-time dependent care [6]. There is currently no approved therapy specific for the treatment of Angelman syndrome.

The maternal-specific inheritance pattern of Angelman syndrome is due to genomic imprinting of *UBE3A* in neurons of the central nervous system (CNS), a naturally occurring phenomenon in which the maternal allele is expressed while the paternal allele is not [7, 8]. The paternal *UBE3A* allele is repressed by the *UBE3A* antisense (*UBE3A-AS*) transcript, which is not a gene per se but rather represents the distal end of the Small Nucleolar RNA Host gene 14 (*SNHG14*) [9]. *SNHG14* — also known as the *IC*-*SNURF-SNRPN* or *U-UBE3A-ATS* — is an imprinted, paternally expressed transcript located downstream and in the opposite orientation of the *UBE3A* gene. It is a structurally complex transcription unit that produces a massive polycistronic transcript (>600 kb) comprised of the *SNURF*/*SNRPN* bicistronic messenger RNAs (mRNA), a long non-coding RNA (lncRNA) host gene that produces an array of orphan C/D box small nucleolar RNAs [*SNORD107*, *SNORD64*, *SNORD109A*, *SNORD116, SNORD115*, and *SNORD109B*], and the *UBE3A-AS* transcript (for review, see Cavaille *et al.* [10]). Interestingly, paternal-derived deletions of *SNHG14* that encompass the *SNORD109A* and *SNORD116* genes cause another neurodevelopmental disorder, Prader-Willi syndrome (PWS), which is characterized by morbid obesity, hypotonia, and mild intellectual disability [11].

One of the main therapeutic strategies currently being explored for Angelman syndrome involves restoring UBE3A protein in the CNS by inhibiting the *UBE3A-AS* transcript to reactivate expression of the paternal *UBE3A* allele [12, 13]. Several studies have shown that this is possible and that reinstating *Ube3a* expression improves the neurological deficits in mouse models of Angelman syndrome, demonstrating the feasibility and potential clinical benefit of a disease-modifying strategy for Angelman syndrome patients [14–20]. Antisense oligonucleotides (ASOs) are an ideal therapeutic modality to inhibit the expression of the *UBE3A-AS* transcript, as they are highly specific and can induce transcriptional termination of actively transcribed genes [21–23]. They can also be administered to patients by lumbar intrathecal injection, where they broadly distribute throughout the CNS [24]. Using a high-throughput screening approach, Meng *et al*. [16] developed ASOs targeting the mouse *Ube3a-AS* transcript and demonstrated cognitive improvement in a mouse model of Angelman syndrome. It is unclear, though, if these ASOs will translate into clinical practice because of the substantial molecular evolutionary divergence known to exist between humans and mice [25].

Here, we describe the development of human-specific ASOs for the treatment of Angelman syndrome patients. We first examined how the human *UBE3A-AS* transcript and mouse *Ube3a-AS* transcript are regulated in the adult brain. We then used a multi-faceted comparative approach to identify an ASO target region that would precisely inhibit transcription of the *UBE3A-AS* transcript. Lastly, we developed and optimized ASOs and tested their pharmacological properties in cultured human neurons and in a non-human primate model using a clinically relevant route of administration.

## RESULTS

### The UBE3A-AS transcript is a spliced, polyadenylated transcript regulated as part of the host gene

The regulation of the *SNHG14* transcription unit and the *UBE3A* gene is complex (**Fig. 1A**). It is currently thought that transcription of *SNHG14* initiates on the paternal allele at promoter(s) upstream of *SNURF*/*SNRPN* and terminates at one of two sites downstream. In non-CNS tissues, transcription ends between the *SNORD116* and *SNORD115* gene arrays, whereas in CNS neurons, it extends downstream to express the *SNORD115* and *SNORD109B* genes and the *UBE3A-AS* transcript. The antisense transcription of the *UBE3A-AS* transcript is then thought to repress the expression of the paternal *UBE3A* allele by transcriptional interference. Although poorly understood, the nascent *SNHG14* transcript is co- and post-transcriptionally processed into a complex population of RNAs. Whether the *UBE3A-AS* transcript is a processed transcript or simply reflects transcriptional noise is debated [9, 26–29]. To address this question, we performed strand-specific RNA-sequencing on total RNA (toRNA-seq) and polyA-enriched RNA (mRNA-seq) isolated from the cerebral cortex of a control individual (n = 1) and then examined the aligned sequence reads and assembled transcripts. We also analyzed publicly available polyA-seq data to identify the sites at which the *SNHG14* transcript is polyadenylated in the human brain.

**Figure 1.**
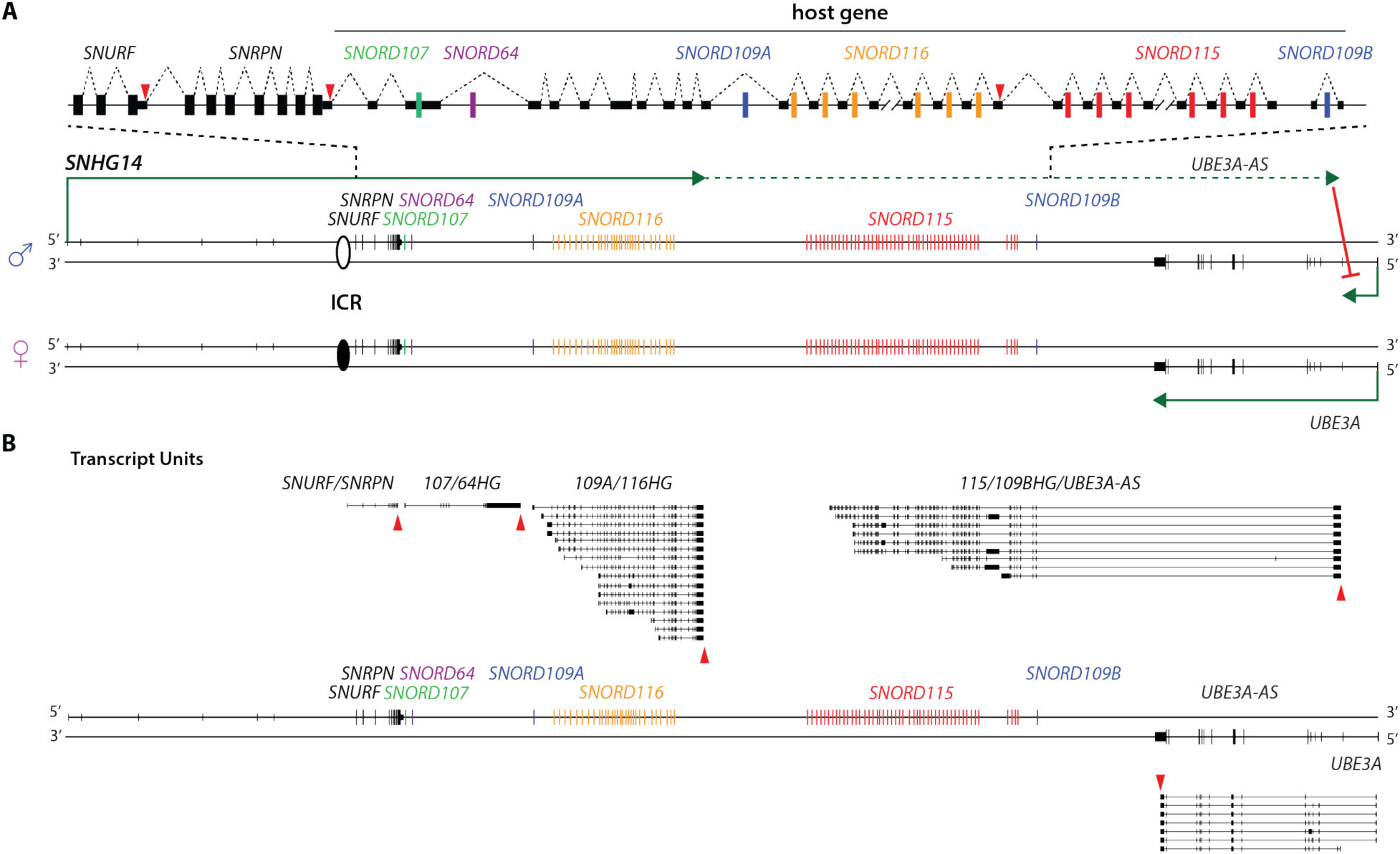
The *UBE3A-AS* transcript is a spliced, polyadenylated transcript regulated as part of the host gene. (**A**) Schematic illustrating the imprinted and tissue-specific regulation of the human *SNHG14* transcription unit and *UBE3A*. The *SNHG14* polycistronic transcript encodes the *SNURF*/*SNRPN* protein-coding mRNAs, *SNORD107*, *SNORD64*, *SNORD109A*, *SNORD116* (29 copies), *SNORD115* (48 copies), *SNORD109B*, and the *UBE3A-AS* transcript. Differential methylation of an imprinting control region (ICR) regulates the paternal-specific expression of *SNHG14*. The white and black ovals represent unmethylated (active) and methylated (inactive), respectively. The proximal end of *SNHG14* is transcribed in all tissues (solid line), whereas the distal end is only transcribed in CNS neurons (dashed line), which represses the paternal *UBE3A* allele. The nascent transcript is processed by splicing (dashed lines) and the 3’-end cleavage and polyadenylation (red triangles) to generate the different RNAs. The host gene consists of non-coding exons flanking an intron-embedded *SNORD* RNA. Splicing of the exons and processing of the intron generates mature *SNORD* RNAs. (**B**) Analysis of strand-specific mRNA-seq and polyA-seq indicates that the *SNHG14* nascent transcript is processed into separate transcript units by 3’ cleavage and polyadenylation, indicated by red triangles. The *UBE3A-AS* transcript is regulated as part of the distal host gene, where it is alternatively spliced and polyadenylated. The black boxes and lines represent exons and introns, respectively.

Analysis of the data sets indicated that *SNHG14* is transcribed as a single, long transcript that is subject to alternative splicing and 3’-end polyadenylation and cleavage to produce separate transcript units corresponding to the *SNURF*/*SNRPN* mRNA, the *SNORD107*/*SNORD64* host gene (*107*/*64HG*), the *SNORD109A*/*SNORD116* host gene (*109A*/*116HG*), and the *SNORD115*/*SNORD109B* host gene and *UBE3A-AS* transcript [*115*/*109BHG*/*UBE3A-AS*) (**Fig. 1B and fig. S1A and S1B**)]. Analysis of the *115*/*109BHG*/*UBE3A-AS* transcript unit showed that the host gene transcript splices into exons of the *UBE3A-AS* transcript, which itself is alternatively spliced and polyadenylated (**Fig. 1B and fig. S1B**). Closer inspection revealed that the *UBE3A-AS* transcript is 1) comprised mainly of intronic sequence, with the major splice variant having an extremely large intron (140 kb) that spans from the distal host gene exon of *SNORD109B* to the 3’-terminal exon at the 5’-end of *UBE3A*, 2) co-transcriptionally spliced, and 3) mostly polyadenylated (**fig. S1C**). The splicing pattern and polyadenylation of the *115*/*109BHG*/*UBE3A-AS* transcript were verified using long-read Oxford Nanopore sequencing (**fig. S1D**). Altogether, these findings show that *SNHG14* is processed into separate transcript units by 3’-polyadenylation and cleavage, consistent with and expanding on prior findings [30], and that the *UBE3A-AS* transcript is a spliced, polyadenylated component of the host gene.

### The mouse Ube3a-AS transcript is regulated as part of the Snord host gene but has diverged substantially from human and other mammalian orthologs

The organization of the mouse *Snhg14* transcription unit is conserved, to a large extent, with humans; however, it contains additional copies of the *Snord116* and *Snord115* genes and lacks orthologs of the *SNORD109A*/*B* genes (**fig. S2A**) [31, 32]. Therefore, we performed comparable transcriptional profiling to better understand the regulation of the *Ube3a-AS* transcript and determine the extent to which it is conserved with the human *UBE3A-AS* transcript. Our analyses indicated that the mouse *Snhg14* transcript is also processed into different transcript units and that the *Ube3a-AS* transcript is also regulated as part of the host gene (**fig. S2B and S2C**). In contrast to the human *UBE3A-AS* transcript, however, we found that the mouse *Ube3a-AS* transcript is extensively spliced and has alternate polyadenylation sites in and upstream of *Ube3a* (**fig. S2C**). Furthermore, we found no sequence or structural homology between the human and mouse *UBE3A-AS*/*Ube3a-AS* transcripts. Taken together, these findings show that, although the mouse *Ube3a-AS* transcript is regulated as part of the host gene and polyadenylated, its sequence and genetic structure have diverged substantially. These results further indicate that the sequence-specific therapies targeting the mouse *Ube3a-AS* transcript are not likely to translate to humans.

### The start of the UBE3A-AS transcript is genetically unique and evolutionarily conserved

Based on our findings, we next sought to identify an ASO target region that would specifically and efficiently inhibit transcription of the *UBE3A-AS* transcript prior to the 3’-end of *UBE3A*. The highly repetitive nature of the host gene and the general lack of knowledge of the specific regulatory sequences in the region posed several challenges. We thus examined the sequence composition and genetic architecture of *SNHG14* to identify a potential target region with the following characteristics: 1) unique sequences in the distal transcript unit and not duplicated elsewhere in *SNHG14*, 2) a region with evidence of regulatory potential, and 3) a region conserved between humans and a suitable animal model to enable *in vivo* testing of candidate ASOs.

To identify repeated structures in the *SNHG14* transcription unit, we first examined dot plots constructed from self-alignments. We discovered several crucial features in the region. First, we found that the *SNORD109B* host gene exons are duplicated upstream at *SNORD109A* (**Fig. 2A and fig. S3A - C**), and thus ASOs targeting the region spanning *SNORD109B* would likely have off-target effects at *SNORD109A*, potentially dysregulating the Prader-Willi syndrome critical region. Second, we made the novel observation that the *SNORD115* gene array is partitioned into two distinct clusters, one that is highly repetitive (Cluster 1) and one that is a single copy (Cluster 2) (**Fig. 2B and fig. S4A**). We extended our analysis of these two clusters and found that the host gene exons in Cluster 1 (E-1 and E-2 exons) share high sequence identity across the repeated units, and thus ASOs targeting this region would likely have multiple off-target effects, potentially dysregulating the entire *SNORD115* gene array (**Fig. 2B and fig. S5A**)]. In contrast, we found that Cluster 2 is organized as five different host gene exons (E-3 exons) that span a cluster of *SNORD115* pseudogenes (**Fig. 2C and fig. S5B and fig. S6A**). Although similar in size, the E-3 exons shared little, if any, sequence identity to each other, to the exons in Cluster 1, or to any other sequences in *SNHG14*. Importantly, this region represents the structural start of the *UBE3A-AS* transcript. Finally, we found that the two clusters have strikingly different sequence compositions (GC content and repetitive elements) and are separated by an unusually large intron, creating a boundary-like element separating the functional *SNORD115* genes in Cluster 1 from Cluster 2, *SNORD109B*, and the *UBE3A-AS* transcript (**Fig. 2D and fig. S7A)**. Altogether, these findings show that Cluster 2 is genetically unique, and thus ASOs targeting this region would be specific to the start of the *UBE3A-AS* transcript.

**Figure 2.**
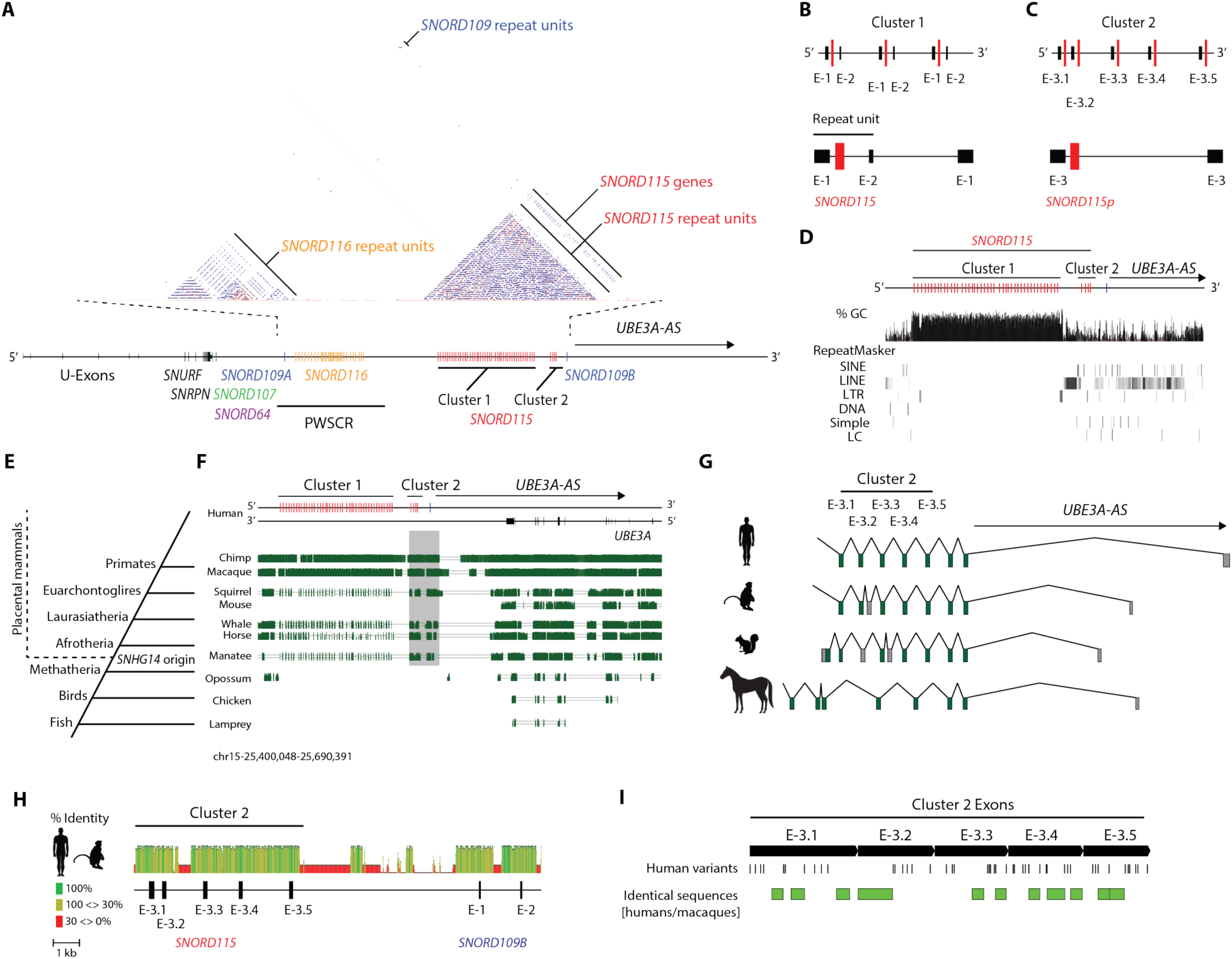
The start of the *UBE3A-AS* transcript is genetically unique and evolutionarily conserved. (**A**) A dot plot constructed from a self-alignment of the *SNHG14* transcription unit shows that the *SNORD109*, *SNORD116*, and *SNORD115* genes are organized as clusters of different repeated units. The *SNORD109A* and *SNORD109B* genes and host gene exons are duplicated at the proximal and distal ends of the region. The *SNORD115* gene array is partitioned into two distinct clusters: Cluster 1 and Cluster 2. The sequence of Cluster 2 is divergent from Cluster 1, except for the *SNORD115-48* gene, which shares homology with the *SNORD115* genes in Cluster 1. (**B - C**) Schematic representations of the repeat units in Cluster 1 and Cluster 2 (drawn to scale). Cluster 1 is organized as repeated units of two different host gene exons (E-1 and E-2) flanking an intron-embedded *SNORD115* gene, whereas Cluster 2 is organized as five distinct exons (E-3) flanking a *SNORD115* pseudogene (*SNORD115p*), except for *SNORD115-48*. The E-3 exons are similar in size but have divergent sequences. (**D**) Schematic of the guanine-cytosine content (%GC) and distribution of repetitive elements in Cluster 1 and Cluster 2. Abbreviations: SINE, small interspersed nuclear elements; LINE, long interspersed nuclear elements; LTR, long terminal repeat; DNA, DNA repeats; Simple, simple repeats; LC, low copy repeats. (**E**) Phylogenetic tree illustrating the evolution of the *SNHG14* transcription unit in a common ancestor of placental mammals approximately 105 to 180 million years ago (MYA). (**F**) Multiz genome alignments between humans and the orthologous regions in distantly related animals show long stretches of conserved sequence blocks spanning Cluster 2 and *SNORD109B* (grey highlight). Vertical green lines depict conserved sequences. (**G**) Schematic illustrating conservation of the *UBE3A-AS* transcript in human, cynomolgus macaque, squirrel, and horse. The E-1 and E-2 exons (not shown) are not conserved. The E-3 and *SNORD109B* exons are conserved and represent the structural start of the *UBE3A-AS* transcript. Boxes and lines represent exons and introns, respectively. The green and grey boxes represent the conserved and divergent exons, respectively. (**H**) Schematic showing percent (%) identity between humans and cynomolgus macaques in the region spanning Cluster 2 and *SNORD109B.* (**I**) Schematic showing identical sequences across humans and cynomolgus macaque.

To identify potential regulatory elements, we examined the distal end of *SNHG14* for highly conserved sequences across placental mammals, as evolutionarily constrained sequences have been shown to be indicative of regulatory potential [33]. It is thought that imprinting of *UBE3A* was acquired in a common ancestor of placental mammals over 100 million years ago, after the assembly of the *SNHG14* transcription unit downstream of *UBE3A* and the evolution of the *UBE3A-AS* transcript (**Fig. 2E**) [34, 35]. Based on these prior findings and our results showing that the *UBE3A-AS* transcript is regulated as part of the host gene, we hypothesized that evolutionarily conserved sequences at the distal end of *SNHG14* might be involved in regulating the *UBE3A-AS* transcript. To test this, we examined Multiz genome alignments between humans and the orthologous regions in distantly related animals.

We detected long stretches of highly conserved sequence blocks spanning Cluster 2 and *SNORD109B*, which contrasted with much shorter stretches of conserved sequences upstream in Cluster 1 and the lack of sequence conservation downstream, between *SNORD109B* and the 3’- end of *UBE3A* (**Fig. 2F**). Closer inspection showed that the conserved sequences in Cluster 1 primarily corresponded to the intron-embedded *SNORD115* genes, whereas the conserved sequences in Cluster 2 corresponded to the host gene introns and exons (**fig. S8A and S8B**), indicating that the two clusters have been subjected to different evolutionary constraints. Examination of orthologous *SNHG14*/*UBE3A-AS* transcripts from cynomolgus macaques, squirrels, and horses further showed that only the E-3 and *SNORD109B* exons are conserved and that these exons demarcate the start of the *UBE3A-AS* transcript in each species (**Fig. 2G**). Additional analyses revealed stretches of identical sequences between humans, apes, and Old-World monkeys in Cluster 2 (**Fig. 2H and 2I**), indicating that these sequences have remained unchanged for at least 29 million years. Collectively, these findings indicate that both the sequence and transcriptional regulation of the E-3 and *SNORD109B* exons have been remarkably conserved throughout placental mammalian evolution — although uniquely lost in mouse — suggesting the acquisition of this region was critical in establishing and regulating the expression of the *UBE3A-AS* transcript. These results further show that non-human primates (NHP) are a suitable animal model to test human-specific ASOs targeting sequences in Cluster 2.

### ASOs targeting exons in Cluster 2 reverse imprinting of UBE3A in cultured human neurons

Based on our findings, we next sought to determine whether gapmer ASOs targeting Cluster 2 would terminate transcription of the *UBE3A-AS* transcript and reactivate expression of the paternal *UBE3A* allele. We targeted the E-3 exons as our findings indicated that they are co-transcriptionally spliced, and thus the introns would be excised from the nascent transcript during transcription. We designed six human-specific ASOs, including three ASOs targeting identical sequences between humans and cynomolgus macaques, a commonly used NHP model (**Fig. 3A**, **fig. S9A, S9B, and Table S1**). For our initial screening, we designed ASOs, 20 nucleotides in length (20-mer), with phosphorothioate (PS) linkages and 2’-O-methyl (O) modified ribonucleosides (**Fig. 3B and 3C**). To assess the pharmacology of the ASOs, we treated GABAergic neurons derived from a human induced pluripotent stem cell line with the ASOs. We then quantified the steady-state RNA levels of the *UBE3A-AS* transcript using quantitative RT-PCR (qRT-PCR). For comparison, we treated the neurons with Topotecan, a chemotherapeutic agent that potently inhibits *UBE3A*-AS expression [36, 37].

**Figure 3.**
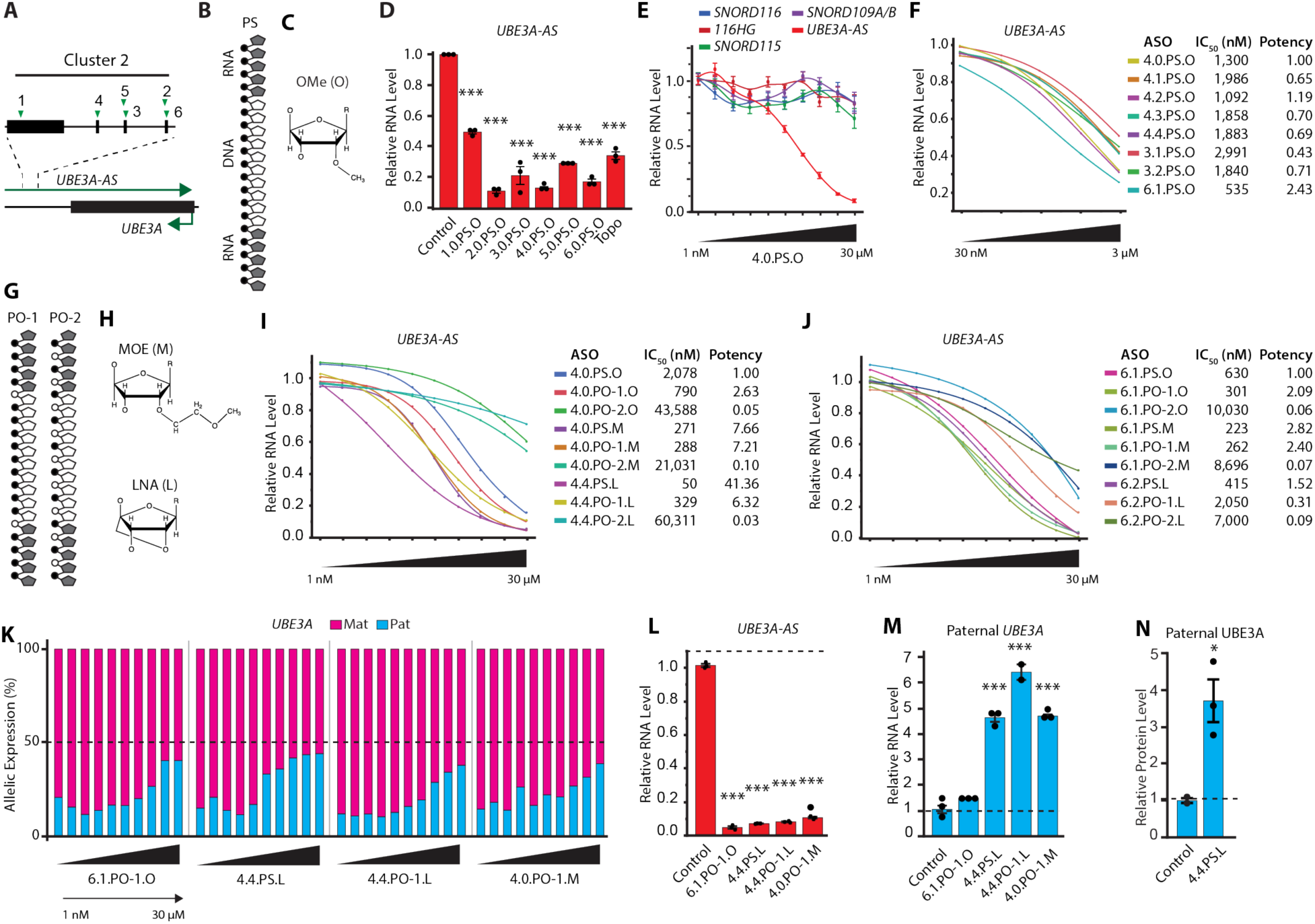
ASOs targeting exons in Cluster 2 reverse imprinting of *UBE3A* in human neurons. (**A**) Schematic showing the ASO target sites (green triangles) in Cluster 2. Black boxes and lines represent exons and introns, respectively. (**B**) Schematic of gapmer ASO with phosphorothioate (PS) linkages. The modified ribonucleosides and deoxynucleosides are depicted by grey and white pentagons, respectively. The PS linkage is depicted by a black circle. (**C**) Schematic of 2’-O- methyl modified ribonucleoside [OMe (O)]. (**D**) *UBE3A-AS* RNA levels in GABAergic neurons treated with 1.0.PS.O, 2.0.PS.O, 3.0.PS.O, 4.0.PS.O, 5.0.PS.O, 6.0.PS.O (10 µM, n = 3 wells/treatment) and Topotecan (Topo, 1 µM, n = 2 wells), normalized to control (n = 3). Data presented as mean ± standard error of mean (SEM); Mixed effect linear regression followed by Tukey HSD test, \**P* < 0.05, \*\**P* < 0.01, \*\*\**P* < 0.0001. Neurons were treated at 14 days in vitro (DIV) and harvested for analysis at 20 DIV. (**E**) The RNA levels of the *SNORD116*, *SNORD115*, and *SNORD109A*/*B* genes, the distal end of the *SNORD116* host gene (*116HG*), and the *UBE3A- AS* transcript in GABAergic neurons treated with a 10-point ½ log dose curve (1 nM, 3 nM, 10 nM, 30 nM, 100 nM, 300 nM, 1 µM, 3 µM, 10 µM, and 30 µM) of 4.0.PS.O (n = 3 wells/treatment), normalized to the controls. Data presented as mean ± SEM. (**F**) *UBE3A-AS* RNA levels in GABAergic neurons treated with a 5-point ½ log dose curve (30 nM, 100 nM, 300 nM, 1 µM, and 3 µM) of 3.1.PS.O, 3.2.PS.O, 4.0.PS.O, 4.1.PS.O, 4.2.PS.O, 4.3.PS.O, 4.4.PS.O, and 6.1.PS.O (n = 2 wells/treatment), normalized to control (n = 5). Data presented as fitted logistic regression curves. IC_50_ values and Relative Potency were estimated using a 3-parameter-logistic regression model and parallelism test, respectively. (**G**) Schematic of chemically optimized gapmer ASOs with different backbone designs (PO-1 and PO-2). The modified ribonucleosides and deoxynucleosides are depicted by grey and white pentagons, respectively. The PS and phosphodiester (PO) linkages are depicted by black and white circles, respectively. (**H**) Schematic of modified 2’-methoxyethyl [MOE (M)] and locked nucleic acid [LNA (L)] ribonucleosides. (**I - J**) *UBE3A-AS* RNA levels in GABAergic neurons treated with a 10-point ½ log dose curve of the chemically modified ASOs (n = 2 wells/treatment), normalized to control (n = 4). Data presented as mean ± SEM. Neurons were treated at 18 DIV and harvested for analysis at 24 DIV. IC_50_ values and Relative Potency were estimated using a 4-parameter-logistic regression model and parallelism test, respectively. (**K**) *UBE3A* allelic ratios in GABAergic neurons treated with a 10-point ½ log dose curve of 6.1.PO-1.O, 4.4.PS.L, 4.4.PO-1.L, and 4.0.PO-1.M (n = 3 wells/treatment). The dashed line represents the equal (50:50) expression of the maternal and paternal *UBE3A* alleles. The maternal and paternal alleles are depicted in pink and blue, respectively. Neurons were treated at 23 DIV and harvested for analysis at 29 DIV. (**L - M**) *UBE3A-AS* and paternal *UBE3A* RNA levels in Angelman syndrome neurons treated with 6.1.PO- 1.O, 4.4.PS.L, 4.4.PO-1.L, and 4.0.PO-1.M (30 µM, n = 3 wells/treatment), normalized to control (n = 3, dashed line). Data presented as mean ± SEM; Mixed effect linear regression followed by Dunnet-HSU test, \*\*\**P* < 0.0001. (**N**) Paternal UBE3A protein levels in Angelman syndrome neurons treated with 4.4.PS.L (30 µM, n = 3 wells/treatment), normalized to control (n = 2, dashed line). Neurons were treated at 66 DIV and harvested for analysis at 72 DIV. Data presented as mean ± SEM; Mixed effect linear regression followed by Dunnet-HSU test, \**P* < 0.05.

The ASOs significantly reduced *UBE3A-AS* RNA levels, with three ASOs achieving greater than 85% reduction (**Fig. 3D**). Additional studies showed that the ASO treatment specifically reduced *UBE3A-AS* RNA levels in a dose-dependent manner but not the RNA levels of the *SNORD116*, *SNORD115*, or *SNORD109A*/*B* genes nor the *SNORD116* host gene (**Fig. 3E**). We then optimized the target sequences and chemistry of the three conserved ASOs (3.0.PS.O, 4.0.PS.O, and 6.0.PS.O), which were chosen as lead candidates due to their pharmacological properties and conservation with cynomolgus macaques. To optimize the target sequences, we designed seven additional ASOs targeting slightly different sequences within the conserved regions of the exons (**Table S2 and fig. S10A**). Analysis of the ASOs in the GABAergic neurons revealed two ASOs (4.0.PS.O and 6.1.PS.O) with slightly improved pharmacological properties (**Fig. 3F and Table S3**). To optimize the chemistry of 4.0.PS.O and 6.1.PS.O, we designed 16 additional ASOs targeting the same sequence but with two different backbone designs [mixed phosphorothioate and phosphodiester (PO-1 and PO-2)] and two different chemically modified ribonucleosides [2’-methoxyethyl (M) and locked nucleic acid (L) **(Fig. 3G and 3H, Table S4, and fig. S10B and S10C)**]. The size of the LNA-modified ASOs was reduced to account for the increased affinity associated with this modification [38]. Analysis of the ASOs in the GABAergic neurons revealed nine ASOs more potent than 4.0.PS.O and 6.1.PS.O, with each ASO having half-maximal inhibitory potencies (IC_50_) in the nanomolar range and achieving greater than 90% reduction of *UBE3A-AS* RNA (**Fig. 3I, 3J, and Table S5**).

To determine whether the ASOs reactivated paternal *UBE3A* expression, we used a heterozygous single nucleotide polymorphism (SNP) identified in the *UBE3A* gene of the GABAergic neurons. Using an allele-specific digital droplet RT-PCR (ddRT-PCR) assay, we found that the inferred maternal *UBE3A* allele was preferentially expressed relative to the paternal *UBE3A* allele [maternal:paternal allelic ratio = 87:13] in the neurons, similar to the allelic expression of *UBE3A* in the human brain [8]. Additional experiments using four of the most potent ASOs (6.1.PO-1.O, 4.4.PS.L, 4.4.PO-1.L, and 4.0.PO-1.M) showed that each ASO increased expression of the paternal *UBE3A* allele in a dose-dependent manner, with the highest concentration achieving almost complete biallelic expression (**Fig. 3K and Table S6**). To confirm these findings, we tested the ASOs in an iPSC neuronal cell line derived from an Angelman syndrome patient harboring a deletion of the maternal 15q11-q13 region. As expected, the ASOs significantly reduced *UBE3A-AS* RNA levels and increased paternal *UBE3A* RNA and UBE3A protein levels (**Fig. 3L, 3M, and 3N**). Collectively, these findings show that ASOs targeting the E-3 exons in Cluster 2 specifically repress the *UBE3A-AS* transcript and reactivate the expression of the paternal *UBE3A* allele in cultured human neurons.

### ASOs reduce UBE3A-AS and increase paternal UBE3A RNA levels in the CNS of non-human primates

Our ASO design allowed us to assess the pharmacological properties of the lead candidates in an NHP model using lumbar intrathecal injection, the intended route of administration in Angelman syndrome patients. To assess whether the regulation of the cynomolgus macaque *SNHG14*/*UBE3A-AS* transcript is conserved with humans, we generated and analyzed mRNA-seq data as described above for human and mouse. Our analysis indicated that the cynomolgus macaque *SNHG14* transcript is regulated like the human *SNHG14* transcript, in particular the *UBE3A-AS* transcript (**fig. S11A - C**).

To assess the *in vivo* pharmacological properties of the ASOs, we tested four different ASOs, representing three different target sequences, two backbone designs, and three RNA modifications (6.1.PO-1.O, 4.4.PS.L, 4.4.PO-1.L, and 4.0.PO-1.M). The ASOs were administered to the animals by lumbar intrathecal injection using three different study designs consisting of single or repeat doses with different concentrations of ASO (Studies 1 – 4). At the end of each study, we used qRT-PCR to quantify the steady-state RNA levels of the *UBE3A-AS* transcript in eleven CNS regions (**fig. S12A**). Our analysis showed that the ASOs reduced *UBE3A-AS* RNA levels in different CNS regions, with the most significant effects occurring in the lumbar spinal cord, motor cortex, frontal cortex, and hippocampus (**Fig. 4A – D and fig. S12B - E**). Two ASOs (4.4.PS.L and 4.4.PO-1.L) significantly reduced *UBE3A-AS* RNA levels in a dose-dependent manner, achieving greater than 85% reduction in several regions (**Fig. 4D and fig. S12E**). Overall, the effect of ASO treatment was different between ASOs, CNS regions, and dosing regimens.

**Figure 4.**
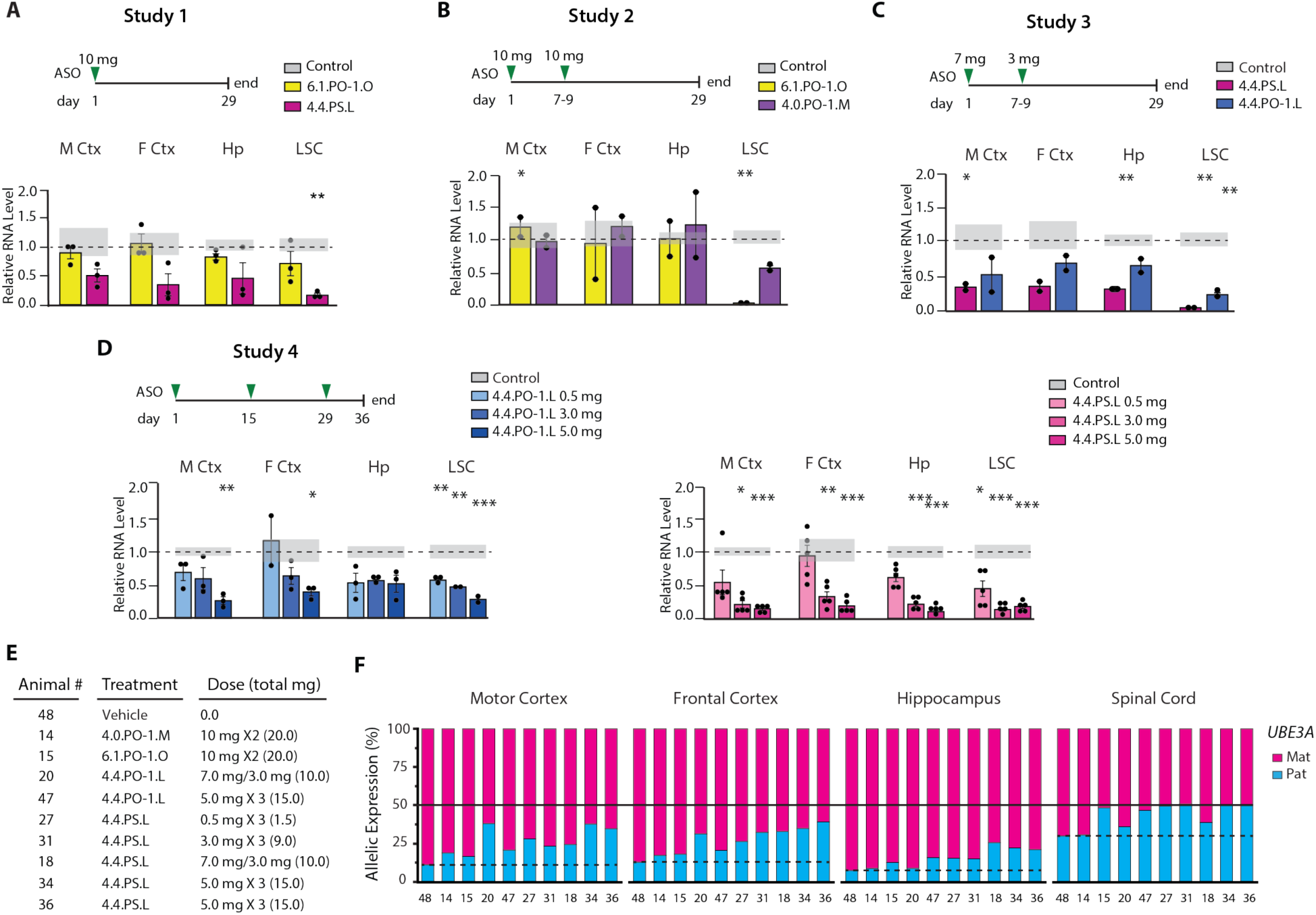
ASO treatment reduces *UBE3A-AS* RNA and reactivates paternal *UBE3A* expression in the cynomolgus macaque CNS. (**A**) *UBE3A-AS* RNA levels after a single injection of 6.1.PO-1.O (10 mg, n = 3) and 4.4.PS.L (10 mg, n = 3), normalized to control [n = 5 (dashed line, mean; grey highlight, SEM)]. Data presented as mean ± SEM; Mixed effect linear regression followed by Dunnett’s test, \**P* < 0.05, \*\**P* < 0.01, \*\*\**P* < 0.0001. (**B**) *UBE3A-AS* RNA levels after two injections of 6.1.PO-1.O [10 mg, 10 mg (n = 3)] and 4.0.PO-1.M [10 mg, 10 mg (n = 3)], normalized to control [n = 6 (dashed line, mean; grey highlight, SEM)]. Data presented as mean ± SEM; Mixed effect linear regression followed by Dunnett’s test, \**P* < 0.05, \*\**P* < 0.01, \*\*\**P* < 0.0001. (**C**) *UBE3A-AS* RNA levels after two injections of 4.4.PS.L [7 mg, 3 mg (n = 2)] and 4.4.PO-1.L [7 mg, 3 mg (n = 2)], normalized to control [n = 6 (dashed line, mean; grey highlight, SEM)]. Data presented as mean ± SEM; Mixed effect linear regression followed by Dunnett’s test, \**P* < 0.05, \*\**P* < 0.01, \*\*\**P* < 0.0001. (**D**) *UBE3A-AS* RNA levels after three injections of 4.4.PO-1.L or 4.4.PS.L [0.5 mg, 3.0 mg, and 5.0 mg (n = 3/dose)] normalized to control [n = 3 (dashed line, mean; grey highlight, SEM)]. Data presented as means ± SEM; Mixed effect linear regression followed by Dunnett’s test, \**P* < 0.05, \*\**P* < 0.01, \*\*\**P* < 0.0001. (**E**) Animals harboring a single nucleotide variant in the *UBE3A* gene. (**F**) Parental *UBE3A* allelic expression in individual animals after ASO treatment. The maternal and paternal alleles are depicted in pink and blue, respectively. Paternal allelic expression of the control animal is depicted by a dashed line. Biallelic expression of *UBE3A* (50:50) is depicted by a solid line.

To examine reactivation of the paternal *UBE3A* allele, we used a heterozygous single nucleotide variant (SNV) identified in the *UBE3A* gene of a subset of the animals (n = 10, **Fig. 4E and Table S7**). Using an allele-specific ddRT-PCR assay, we analyzed the allelic expression patterns of *UBE3A* in the motor cortex, frontal cortex, hippocampus, and lumbar spinal cord. Our analysis showed that paternal *UBE3A* expression increased in the motor cortex, frontal cortex, and spinal cord of several animals, with complete biallelic expression achieved in the spinal cord of six animals (**Fig. 4F and Table S8 and S9**). Altogether, these findings demonstrate that human-specific ASOs administered to cynomolgus macaques by lumbar intrathecal injection repress the *UBE3A-AS* transcript and increase paternal *UBE3A* expression in several regions of the CNS.

## DISCUSSION

Reactivation of the paternal *UBE3A* allele is currently considered one of the most promising therapeutic options for treating Angelman syndrome. Using a multi-faceted approach to understand the regulation, genetic structure, and conservation of the *SNHG14* transcription unit, we identified a highly conserved region that we believe represents the ancestral start site of the *UBE3A-AS* transcript. We demonstrate that the ASOs targeting this region specifically and efficiently repress the *UBE3A-AS* transcript and increase paternal *UBE3A* expression in cultured neurons and in the CNS of cynomolgus macaques. Results from this study have advanced the first molecular therapy for Angelman syndrome into clinical development (ClinicalTrials.gov, NCT04259281).

Given the widespread use of mouse models in Angelman syndrome research and the recent reports involving ASOs and CRISPR/Cas9 to reactivate paternal *Ube3a* expression [16, 17, 39], we thought it was essential to assess the conservation of the mouse *Ube3a-AS* transcript with the human *UBE3A-AS* transcript. Our findings that the mouse *Ube3a-AS* transcript has diverged substantially from humans indicate that the sequence-based therapies targeting this region in mice will likely not translate to human patients. Furthermore, our finding that this region has diverged from all other placental mammals, including other rodents (e.g., squirrels), is consistent with the accelerated rate of evolution observed in murid species [40, 41]. It should also serve as a cautionary tale, illustrating the large extent to which the mouse genome may diverge from other placental mammals, particularly at functionally relevant regions of the human genome.

Our discovery that the *SNORD115* gene array is partitioned into two genetically and structurally distinct clusters is important for several reasons. First, based on the results from prior studies examining the evolutionary origin of the *SNORD115* genes [42] and our comparative genomics studies, we propose that Cluster 2 represents the ancestral origin of the *UBE3A-AS* transcript. The region has been remarkably conserved for a non-coding RNA— at least 95 million years based on the evolutionary divergence between humans and horses — and has retained its regulatory potential, as evidenced by the conserved splicing pattern of the E-3 and *SNORD109B* exons as part of the *UBE3A-AS* transcript. Though additional phylogenetic studies are needed, we hypothesize that the ancestral host gene exon(s) in Cluster 2 were co-opted to generate the *UBE3A-AS* transcript, resulting in the imprinting of *UBE3A*. Second, the E-3 exons are divergent from each other and unique in the genome, making them ideal sequences to target with ASOs. These unique sequences and the degree to which the exons are conserved across NHPs provided a way to specifically target the *UBE3A-AS* transcript and test human-specific ASOs in a large animal model using a clinically relevant route of administration. Altogether, this approach allowed us to develop and test clinical ASO candidates and infer their pharmacological properties in the CNS.

Overall, our results show that several of the ASOs can precisely inhibit the *UBE3A-AS* transcript and efficiently reactivate the expression of the paternal *UBE3A* allele in cultured neurons. Importantly, we show that lumbar intrathecal administration of several ASOs can reduce *UBE3A-AS* RNA levels and increase paternal *UBE3A* expression in several disease-relevant regions of the cynomolgus macaque CNS. Two ASOs reduced *UBE3A-AS* RNA levels by 60% to 85% in several brain regions and by as much as 90% in the lumbar spinal cord. The reduction in *UBE3A-AS* levels correlated with increased paternal *UBE3A* RNA expression, achieving as much as 59 to 62% of the maternal *UBE3A* allele in the cerebral cortex and complete biallelic expression of *UBE3A* in the spinal cord. Although the therapeutic benefit of an ASO treatment can only be assessed in human patients, we anticipate that this level of paternal *UBE3A* reactivation would likely be clinically beneficial in Angelman patients, given the findings from studies in mouse models of Angelman syndrome and observations made in patients with rare mutations [15, 16, 19, 43–45].

Though we believe the findings from this study are significant, we acknowledge several limitations. First, despite the advantages of using an NHP model to test our investigational ASOs, the presence of a functional maternal *UBE3A* allele confounded our ability to evaluate paternal UBE3A protein levels. Therefore, we were unable to correlate reactivation of paternal *UBE3A* expression with increased paternal UBE3A protein levels, nor could we determine the half-life of paternal UBE3A protein after reactivation. Second, additional studies are needed to assess the distribution and half-life of the ASOs in the CNS. Lastly, the sample size for several of the NHP studies was relatively small, especially in our analysis of paternal *UBE3A* expression, which was limited to animals harboring a heterozygous SNP in the *UBE3A* gene.

In conclusion, we describe the development and characterization of investigational ASOs for the treatment of Angelman syndrome.

## MATERIALS AND METHODS

### Study Design

This study aimed to develop an investigational antisense oligonucleotide (ASO) therapy to treat individuals with Angelman syndrome. The primary objectives of the study were: 1) to understand and compare how the human *SNHG14*/*UBE3A-AS* and mouse *Snhg14*/*Ube3a-AS* transcripts are regulated in the brain, 2) to identify a target region that would specifically and efficiently inhibit the expression of the human *UBE3A-AS* transcript, and 3) to develop and test potential investigational ASOs in cultured human neurons and in a large animal model.

The sample size for the cell culture studies was based on previously published studies testing the effect of ASOs in cultured neurons. Treatments were replicated (n = 2 – 3) for each dose for each experiment and performed on different cell-culture plates to mitigate potential confounding factors. In some instances, the treatments were replicated in subsequent studies. Technical replicates (n = 2 – 3) were performed for the molecular analyses to assess measurement error.

The n values represented the number of wells in a study and were included in the figure legend. The cynomolgus monkey was chosen as the animal model for this study because of its sequence conservation with humans. The sample size for the animal studies was based on prior studies and determined to be the minimum number of animals required to properly characterize the pharmacological effects of the ASO. The studies were also designed such that they did not require an excessive number of animals to accomplish the objectives. When possible, the age, sex, and origin of the animals were matched to mitigate potential confounding factors. The n values represented the number of animals in a study and were included in the figure legend.

The key resources for this study are provided in the Supplementary Materials (**Table S10**).

### Analysis of the human and mouse *SNHG14*/*Snhg14* transcripts

#### RNA-sequencing

Strand-specific RNA-sequencing (RNA-seq) was performed on polyA-enriched RNA (mRNA-seq) and total RNA (ribosomal RNA depleted, toRNA-seq) obtained from an adult human cortex [66-year-old female (540005, Agilent Technologies)] and an adult mouse cortex [eight-week-old (C57BL/6J) male, bred in-house]. The mouse RNA sample was isolated using Qiagen RNAeasy Plus Kit (Qiagen). The RNA concentration was determined using Qubit 4 Fluorometer (ThermoFisher Scientific) using the Qubit BR assay kit (ThermoFisher Scientific), and RNA quality was assessed using an Agilent TapeStation 4200 (Agilent Technologies). The toRNA-seq and mRNA-seq libraries were generated using the Illumina TruSeq Stranded Total RNA kit and Stranded mRNA kit (Illumina) according to the manufacturer’s protocol. 150 base-pair paired-end sequencing was performed using a NovaSeq 6000 (Illumina) at the Texas A&M Institute for Genome Sciences and Society Genomics core (https://genomics.tamu.edu/genomics-core/) and the University of Texas at Arlington North Texas Genome Center (https://northtexasgenomecenter.com).

The raw sequencing reads were processed using bcl2fastq v2.20.0.422 from CASAVA. The resulting FASTQ sequences were examined using FASTQC. The FASTQ sequences were aligned to the mouse (mm9) or human (hg19) reference genome assemblies using Hisat2 (version 2.1.0) [46]. The aligned SAM sequences were then converted to binary BAM sequences, indexed, and sorted using samtools [47]. The aligned sequences were filtered using the view command in samtools to remove non-uniquely aligned reads (quality > 1) [47]. The Watson (5’ - 3’) and Crick (3’ - 5’) strands were separated using the view command in samtools. bedGraph files were generated using the genomecov command in Bedtools [48] and visualized in the UCSC genome browser. The Vertical Range was set to accommodate differences in library sizes.

Transcript assemblies were generated using Stringtie (version 1.3.4.d) [49], with the following settings: -g 10. Single-exon transcripts and transcripts with non-canonical splice-sites were filtered using gffread (GFF utilities, Johns Hopkins University, Center for Computational Biology). Transcript units were defined as overlapping transcripts expressed from the same strand of a transcriptional unit and sharing a common terminal exon. The schematics of the human and mouse *SNHG14*/*Snhg14* transcription units represent the following regions: human (hg19) chr15:25,064,186-25,691,366; mouse (mm9) chr7:66,481,195-66,878,943.

#### PolyA sequencing

The locations of polyadenylation sites were determined using strand-specific polyA-seq data downloaded from the PolyA-seq Track in the UCSC genome browser.

#### Oxford Nanopore RNA-sequencing

Oxford Nanopore MinION long-read sequencing was performed on RNA isolated from an adult human cortex (540005, Agilent Technologies). PolyA-enriched RNA was isolated using the Dynabeads mRNA Direct kit (Life Technologies) following the manufacturer’s protocol. Approximately 250 ng of polyA-enriched RNA was used for input into the Oxford Nanopore Direct cDNA Sequencing library preparation protocol (SQK-DCS108). Long-read sequences were generated on a FLO-Min 106D flow cell (Oxford Nanopore Technologies, Didcot, UK). The raw sequence reads were first error corrected and trimmed using CANU and then aligned to the human reference genome assembly (hg38) using minimap2.

#### Sequence conservation of the human and mouse UBE3A-AS/Ube3a-AS transcripts

Pairwise sequence alignments between the mouse and human *Ube3a-AS*/*UBE3A-AS* transcripts and the corresponding genomic sequences were performed using NCBI BLAST (blastn) [50].

### Sequence analysis of the *SNHG14* transcription unit

#### Self-alignment of the SNHG14 transcription unit

A self-alignment of the *SNHG14* transcription unit [repeat masked, GRCh37 (hg19)] was performed using *Geneious Prime v2021.0.3* and visualized using dot plots based on the *EMBOSS 6.5.7 dottup* tool. A word size of 15 and tile size of 200,000 were selected to decrease uninformative alignments and allow for image export to pdf format. The blue lines indicate alignments shorter than 100 bp, while red lines indicate alignments exceeding 100 bp. Additionally, a self-alignment of the *SNHG14* transcription unit [repeat masked, GRCh37 (hg19)] was performed using NCBI BLAST (discontinuous megablast).

#### Sequence alignments of the host gene and SNORD gene sequences

The sequences of the *SNORD115* host gene were extracted from *SNHG14* (NCBI RefSeq NR_146177) and aligned using the Create Alignment feature in CLC Genomics Workbench (Qiagen Aarhus A/S, version 8.0.1) with the following settings: Gap open cost = 10.0; Gap extension cost = 1.0; End gap cost = Cheap; Alignment mode = Very accurate. The exons were then grouped into clusters based on sequence homology and size. The distance and percent identity of each exon relative to the consensus sequence for its group was estimated using the Comparison feature in CLC Genomics Workbench. The organization of the exons was determined manually using the BLAT feature in the UCSC genome browser.

The *SNORD109A/B* and *SNORD115* gene sequences were downloaded from NCBI RefSeq. Sequence alignments were performed using the Create Alignment feature in CLC Genomics Workbench with the following settings: Gap open cost = 10.0; Gap extension cost = 1.0; End gap cost = Cheap; Alignment mode = Very accurate. *SNORD115* pseudogenes were identified as having degenerate C/D Box sequence motifs [31].

#### Sequence composition analysis

The percentage of guanine and cytosine dinucleotides (GC content) was determined using the Create Sequence Statistics in CLC Genomics Workbench; figures were generated using the GC percent track in the UCSC genome browser. Analysis of repetitive elements was performed using RepeatMasker the Dfam database [51, 52]; figures were generated using the RepeatMasker track in the UCSC genome browser.

### Identification of potential regulatory elements by sequence conservation

#### Multiz genome alignments

Sequence conservation of the distal end of the *SNHG14* transcription unit (hg19 chr15: 25,400,048-25,690,391) was assessed using the Vertebrate Multiz Alignment & Conservation, the Non-primate Placental Mammal Genomes, Chain and Net Alignments, and the Non-placental Vertebrate Genomes, Chain and Net Alignments in the UCSC Genome Browser [53, 54]. Synteny was assessed by identifying reciprocal best alignments via BLAT and the TransMap5 track in the UCSC Genome Browser [55]. NCBI RefSeq genes (curated subset) and GENCODE annotations were obtained from the UCSC Genome Browser [56, 57].

#### Comparative sequence analysis of the SNHG14/UBE3A-AS transcript

*SNHG14*/*UBE3A-AS* transcripts were identified in other species using the Genes and Gene Prediction Tracks in the UCSC Genome Browser and NCBI Genome. *SNHG14* transcripts were identified in cynomolgus macaque [*Macaca fascicularis* (XR_001491839.1)], Squirrel [*Ictidomys tridecemlineatus* (XR_002484396.1)], and Horse [*Equus caballus* (XR_002811332.1)]. Sequence homology between the distal end of the human *SNHG14* transcript and the orthologous transcripts in each species was determined using NCBI BLAST (blastn) with the default settings. Stretches of sequence homology (>20 nucleotides) were then examined in the UCSC genome browser to annotate the conserved sequences. CLUSTAL alignments were generated using the EMBL-EBI search and sequence analysis tools [58].

### Development and optimization of ASOs targeting the *UBE3A-AS* transcript

#### ASO design

Antisense oligonucleotides (ASOs) were designed to target specific regions of the *SNHG14* transcript based on certain criteria. Potential target regions were further filtered based on the presence of known variants present in dbSNP150 and 1000 Genomes Phase 3 Integrated Variant calls [59]. The ASOs were designed using *S*oligo [60, 61]. ASOs with the lowest binding site disruption energy and free binding energy were identified for each target sequence and then inspected for motifs with increased effectiveness. ASOs were further filtered based on accessibility within predicted lowest free energy centroid secondary structure of target sequence generated by *S*oligo. In some instances, secondary structure models were compared using lowest free energy structures generated by RNAfold and Mfold [62]. The ASOs were manufactured by Integrated DNA Technologies, Sigma-Aldrich (St. Louis, MO), Microsynth (Balgach, Switzerland), Eurogentec (Seraing, Belgium), ChemGenes Corporation (Wilmington, MA), and Avecia (Milford, MA).

#### Human induced pluripotent stem cell (iPSC) derived neuronal cultures

GABAergic iPSC-derived neural precursor cells (FujiFilm Cellular Dynamics, R1013) were differentiated into neurons according to the manufactures protocol. Briefly, neural precursor cells were thawed and resuspended in a chemically defined medium and added to sterile-culture plates coated with poly-D-lysine and laminin (ThermoFisher Scientific). The medium was replaced 24 hr after plating, and then one-half of the medium was replaced every 3-5 days afterward.

Angelman syndrome induced pluripotent stem cells (AG1-0 iPSCs) (Kerafast, ECN001) were co-cultured on irradiated murine embryonic fibroblasts in human embryonic stem cell medium [DMEM/F12 (ThermoFisher Scientific), 20% Knockout Serum Replacement (ThermoFisher Scientific), 1X Nonessential amino acids, 2 mM L-glutamine, 7 µl/ml 2-Mercaptoethanol, and 4 µg/ml basic Fibroblast Growth Factor]. For the first passage, AG1-0 cells were passaged according to the product manual for PluriSTEM Human ES/iPS Medium (Millipore Sigma), which is feeder-free and utilizes Dispase II (Millipore Sigma) to dissociate cells. Matrigel™ hESC-qualified Matrix (Corning) was used as an extracellular matrix. At the second passage, the matrix was switched to vitronectin (Millipore Sigma). During subsequent passages, areas of differentiation were manually removed until differentiated cells represented approximately < 5% of the colonies. After four subsequent passages, AG1-0 cells were differentiated using the Millipore ES/iPS Neurogenesis Kit (Millipore Sigma) but lacking vitronectin as an extracellular matrix. The initial passage was performed with EZ-LiFT (Millipore Sigma) to obtain high quality iPS cells. Neural progenitor cells were frozen at stage zero (P_0_) and subsequently thawed in ENSTem-A Neural Expansion Medium (Millipore Sigma) for differentiation. Differentiation was performed on sterile culture plates coated with poly-D-lysine (10 µg/ml) and laminin [10 µg/ml (ThermoFisher Scientific) in differentiation medium (Millipore Sigma) for 10 days of differentiation. In some instances, cells were differentiated in Cellular Dynamics Maintenance Medium (Cellular Dynamics).

### Analysis of gene expression in iPSC-derived neuronal cultures

#### Cell lysis and cDNA synthesis for neuronal culture

Each well was harvested using the Cell-to-CT kit (ThermoFisher Scientific) in a lysate volume of 55 µl. cDNA synthesis was performed with the same kit using an input of 10 µl lysate for a total volume of 50 µl.

#### Quantitative RT-PCR

TaqMan™ quantitative RT-PCR assays were used to assess the steady-state levels of *UBE3A-AS* (Hs01372957_m1), *SNORD115-9, SNORD115-10, SNORD115-12,* and *SNORD115-42* (Hs04275288_gH), *SNORD116HG* (Hs03455409_s1), *SNORD116-11* (Hs04275268_gH), *SNORD109A*/*B* (AP47WVR) in human neurons. *PPIA* (Hs99999904_m1) and *GAPDH* (Hs99999905_m1) were used as internal controls. RNA was isolated from neuronal cultures using the Cell-to-CT kit (ThermoFisher Scientific) in a lysate volume of 55 µl. cDNA synthesis was performed with the same kit using an input of 10 µl lysate for a total volume of 50 µl. TaqMan™ quantitative RT-PCR assays were performed and analyzed as described above.

#### Western blot

Neuronal cultures were lysed in 20 µl of NP40/SDS buffer (1% Nonidet P40/0.01% SDS lysis buffer and protease inhibitors (Roche) per well. The samples were then rotated and centrifuged at 4°C for two 15-min cycles, and the lysate was collected at the end of each cycle. Lysate was prepared for loading for SDS-PAGE by diluting with 4x Laemmli buffer (Bio-Rad). The mixture was heated at 95°C for 10 min and placed on ice. The samples were loaded on 7.5% Mini-PROTEAN® TGX Stain-Free Protein Gels (Bio-Rad). The SDS-PAGE was run at 30V for 40 min and then 100 V for approximately 45 min. The Stain-free gel was then imaged on a UV Transilluminator. The gels were blotted using the Bio-Rad Turbo Transfer System using Trans-Blot Turbo Mini NitroCellulose Transfer Packs (Bio-Rad). The blot was imaged for stain-free loading control after the transfer. The membrane was washed in Tris-buffered saline plus Tween-20 (TBST) and then blocked in 5% milk-TBST for one hr at room temperature. Purified Mouse anti-UBE3A (BD BioSciences, BD611416) was diluted (1:1000) in fresh blocking solution (5% milk-TBST). The membrane was incubated with primary antibody overnight at 4°C. The secondary antibody (HRP – Goat Anti-mouse IgG (H+L), ThermoFisher Scientific) was diluted (1:2000) in 5% milk-TBST. The membrane was incubated with secondary antibody for 1 hr at room temperature. The membrane was developed in Clarity Western ECL Substrates (Bio-Rad) for 5 min, according to the manufacturer’s protocol and then imaged on the Bio-Rad ChemiDoc Imaging System. Data was analyzed using Image Lab 5.2.1. Bands of interest were normalized to the stain-free loading control.

#### Identification of a UBE3A SNP in the GABAergic neurons

RNA-sequencing data from GABAergic neurons was downloaded from the NCBI Sequence Read Archive (SRP056827, SRR1976080, SRR1976078) [63]. Raw FASTQ sequences were processed and aligned to the human reference assembly (hg19) using the methods described above. Visualization of genetic variants was performed using IGV (v2.4.14) [64]. Inspection of the genome sequence of the GABAergic iPSC line revealed a heterozygous, synonymous T/C SNP in exon 6 of *UBE3A* isoform 1 (NM_130838, hg19_chr15:25602024-25602024, rs34670662).

#### Digital droplet RT-PCR analysis of UBE3A allelic expression

The allelic ratios of *UBE3A* in the GABAergic neurons were assessed using digital droplet RT-PCR (ddRT-PCR). A custom TaqMan™ quantitative RT-PCR assay (ANYMTRK) was developed to amplify T (VIC) and C (FAM) alleles of *UBE3A*. Total reaction volume was 25 µl, including 3 µl of cDNA, 12.5 µl of ddPCR SuperMix for Probes (No dUTP) (Bio-Rad), and 1.25 µl of 20X SNP assay. The PCR reaction mix was partitioned into droplets using the QX200 AutoDG Droplet Digital PCR system. RT-PCR amplification was performed using the following conditions: 10 min at 95°C, 40 cycles of 30 sec at 94°C and 1 min at 60°C, and 10 min at 95°C. Droplets were then counted by fluorescence using the QX200 Droplet Reader. Data was analyzed using Quantasoft software (Bio-Rad). The allelic expression was calculated as the fractional abundance of each allele [paternal = paternal/(maternal + paternal); maternal = 1- paternal].

### Analysis of the cynomolgus macaque *SNHG14* transcript

#### RNA-sequencing

The cynomolgus macaque *SNHG14* transcript was assessed using RNA-sequencing data generated from the motor cortex (female, 2.8 years old). RNA isolation, concentration, and quality assessment were performed as described above. PolyA-enriched RNA-sequencing libraries were generated using the Illumina TruSeq Stranded mRNA kit (Illumina), according to the manufacturer’s protocol. 150 base-pair paired-end sequencing was performed using a NovaSeq 6000 (Illumina) at the Texas A&M Institute for Genome Sciences and Society Genomics core and the University of Texas at Arlington North Texas Genome Center as described above. Raw sequencing reads and the resulting FASTQ sequences were processed, analyzed, and visually inspected as described above. The transcript assemblies were generated using Stringtie as described above.

### Pharmacological analysis of ASOs in cynomolgus macaques

#### Animals

Studies involving cynomolgus macaque (*Macaca fascicularis*) were either performed at Charles River Laboratories (CRL, Montreal, Canada) or Northern Biomedical Research Inc (NBR, North Shores, MI). Information for each site is provided below.

The studies performed by CRL were approved by CR MTL Institutional Animal Care and Use Committee (IACUC) before conduct. During the study, the care and use of animals was conducted with guidance from the guidelines of the USA National Research Council and the Canadian Council on Animal Care (CCAC). Each test animal was assigned a unique animal/study number within the population making up the study. The animals were group housed (up to 3/group/cage) or single housed in stainless steel cages with automatic watering systems. Room temperatures were maintained at 20 - 26°C, with humidity maintained at 30 - 70%. The light cycle was 12 hr of light and 12 hr of darkness (except during designated procedures). The animals were fed PMI Certified Primate Diet #5048 twice daily in appropriate amounts for the size and age of the animal. Following surgical procedures, animals received food supplements for 14 days that consisted of bread, fruits and/or vegetables. Psychological/environmental enrichment was provided to the animals as per standard procedure, except during study procedures/activities.

The studies performed by Northern Biomedical Research Inc. complied with all applicable sections of the current version of the Final Rules of the Animal Welfare Act Regulations (9 CFR) and the Guide for the Care and Use of Laboratory Animals, Institute of Laboratory Animal Resources, Commission on Life Sciences, National Research Council, 8th edition. Each test animal was assigned a unique animal/study number within the population making up the study. The animals were housed individually in steel cages with automatic watering systems. Room temperatures were maintained at 72° ± 6°F, with humidity maintained at 50% ± 20%. The light cycle was 12 hr of light and 12 hr of darkness. The animals were fed twenty biscuits of PMI Certified Primate Diet #5048; additional biscuits were given based on the weight and caloric need of the animal. The animals were supplemented with Vitamin C on a weekly basis. Food was withheld at least 12 hr prior to general anesthesia (e.g., dosing or necropsy). Animals were provided with environmental enrichment as per the current NBR Program of Animal Care.

#### Lumbar intrathecal administration of ASOs

4.4.PS.L (10 mg, n = 3): Cynomolgus macaques were of Chinese origin. All animals were anesthetized for the lumbar puncture dosing procedure. The animals were administered dexmedetomidine hydrochloride IM (0.04 mg/kg) for sedation. Approximately 10 min later, an intramuscular injection of ketamine hydrochloride (2.5 mg/kg) was provided to induce further sedation. Once sedated, the animals were intubated and the lumbar area prepared for the spinal tap. The lumbar tap was performed using a Pencan Paed® pencil-point (B Braun, 25-gauge, 50 mm) with the animal in a seated position leaning forward over an acrylic tube to facilitate exposure of the intervertebral space. The puncture was performed at the L5-L6 intervertebral space. After administration, the needle was left in place for approximately 30 sec and then removed. A chlorhexidine-based ointment was placed on the skin over the injection site following removal of the needle. Approximately 0.5 ml of CSF was collected from all animals prior to dosing. The dose volume for each animal was fixed at 1 ml/dose and slowly injected over approximately 1 min. Following dose formulation injection, a saline flush of 0.25 ml was administered. The animals were maintained in prone position, and left in this position for a minimum of 15 min following the flush administration. ASO was reconstituted in 0.9% saline. ASOs were manufactured at Eurogentec (Searaing, Belgium).

6.1.PO-1.O (10 mg, n = 3): Cynomolgus macaques were previously implanted at the lumbar level with an intrathecal catheter connected to a subcutaneous access port as per CRL standard operating procedures. The test and reference item formulation were administered to the appropriate animals by injection via implanted intrathecal catheter and access port on Day 1, or by direct intrathecal injection at the lumbar level under anesthesia. The dose volume for each animal was fixed at 1 ml/dose. Following dose formulation injection, a saline flush of 0.2 ml was administered. The animals were dosed over at least 3 min in prone position and left in this position for a minimum of 15 min following the flush administration. ASOs were manufactured at Avecia.

6.1.PO-1.O [10 mg, 10 mg (n = 3)]: Cynomolgus macaques were previously implanted at the lumbar level with an intrathecal catheter connected to a subcutaneous access port as per CRL standard operating procedures. The test and reference item formulation were administered to the appropriate animals by injection via implanted intrathecal catheter and access port on Day 1, or by direct intrathecal injection at the lumbar level under anesthesia. The dose volume for each animal was fixed at 1 ml/dose. Following dose formulation injection, a saline flush of 0.2 ml was administered. The animals were dosed over at least 3 min in prone position and left in this position for a minimum of 15 min following the flush administration. ASO was reconstituted in aCSF (Harvard Apparatus). ASOs were manufactured at Avecia.

4.0.PO-1.M [10 mg, 10 mg (n = 3)], 4.4.PS.L [7 mg, 3 mg (n = 2)], or 4.4.PO-1.L [7 mg, 3 mg (n = 2)]: Cynomolgus macaques were of Chinese and Cambodian origin. Naïve animals were implanted with a catheter prior to administration of the ASO. Briefly, animals were pretreated with atropine sulfate as a subcutaneous injection at a dose of 0.04 mg/kg. At least 15 min later, an IM dose of 8 mg/kg of ketamine HCl was provided to induce sedation. Animals were intubated and maintained on approximately 1 liter/min of oxygen and 2% isoflurane. Methylprednisolone sodium succinate IV, 30 mg/kg, and flunixin meglumine IM, 2 mg/kg, were administered prior to surgery. A dorsal sagittal incision was made over the lumbar spine and a coextruded polyethylene lined polyurethane catheter (SAI Infusion Technologies) was inserted through a hemilaminectomy of the L4, L5, or L6 lumbar vertebrae. The catheter was advanced to the level of the thoraco-lumbar junction and anchored. The end of the intrathecal catheter was attached to a subcutaneous titanium access port (MINLOVOL, Instech Solomon). The skin was closed with sutures and tissue adhesive. Proper IT-L catheter placement was confirmed with the aid of a myelogram and Iohexol 180. Upon recovery from anesthesia, all animals were provided butorphanol tartrate IM, 0.05 mg/kg, for analgesia and placed on post-surgical antibiotic ceftiofur sodium IM, 5.0 mg/kg, b.i.d. (one injection prior to surgery followed by three injections). A recovery period of at least ten days was provided prior to dosing. The non-naïve animals were previously implanted with a coextruded polyethylene lined polyurethane catheter and subcutaneous titanium MINLOVOL access port. Female cynomolgus monkeys were implanted with intrathecal catheters in the lumbar spine (IT-L) that was used for the dual purpose of dose administration and CSF sample collection. All animals were administered a bolus intrathecal dose of vehicle or ASO through the port/catheter system over approximately 3 min. The dose volume was 1 ml of vehicle or ASO followed by 0.25 ml of aCSF at the same rate to flush the dose from the catheter system. All doses were administered to conscious, unsedated animals at two intervals, approximately one week apart (Day 1 and 8). ASO was reconstituted in aCSF. ASOs were manufactured at ChemGenes Corporation (Wilmington, MA).

4.4.PS.L [3X, 0.5 mg, 3.0 mg, and 5.0 mg (n = 5/dose) and 4.4.PO-1.L [3X, 0.5 mg, 3.0 mg, and 5.0 mg (n = 3/dose): Cynomolgus macaques were of Chinese origin. Animals underwent intrathecal catheterization at the lumbar level as per CR MTL SOPs during the pre-study period. The catheter was connected to a subcutaneous access port to allow for dosing. ASOs were administered to animals by IT lumbar injection via implanted intrathecal catheter and access port on Day 1, 15 and 29. Prior to dosing, the access port site was shaved as necessary, cleaned, and prepared using single use swabstick applicator; manufacturer instructions were followed. The dose volume for each animal was fixed at 1 ml/dose. Following dose formulation injection, a saline flush of 0.2 mL was administered. The animals were dosed over at least 3 min in prone position and left in this position for a minimum of 15 min following the flush administration. ASO was reconstituted in aCSF. ASOs were manufactured at ChemGenes Corporation.

#### Tissue collection

All animals were sedated with 8.0 mg/kg of ketamine HCl IM and maintained on an isoflurane/oxygen mixture and provided with an intravenous bolus of heparin sodium (200 IU/kg). Euthanasia was performed by intravenous injection of sodium pentobarbital followed by exsanguination by incision of the axillary or femoral arteries. At the time of sacrifice, the brain was sectioned at 4-mm coronal sections using a brain matrix. The second slice and every other slice thereafter was used to collect tissue punches for analysis. Four mm tissue punches of lumbar spinal cord (L5 to L6), frontal cortex, globus pallidus, thalamus, temporal lobe, pons, midbrain, motor cortex, hippocampus, medulla, and cerebellum were flash frozen in liquid nitrogen and stored at - 80°C until analysis.

#### RNA extraction and cDNA synthesis

RNA was extracted from half of a 4 mm biopsy punch for each region of the brain using the Qiagen TissueLyser II for tissue disruption with 5mm stainless steel beads (Qiagen). The RNeasy Plus kit (Qiagen) was used for RNA isolation and RNA was eluted in a total of 60 µl sterile H_2_O. RNA concentration was determined using the Qubit 4 Fluorometer (ThermoFisher Scientific) using the Qubit RNA BR assay kit (ThermoFisher Scientific). The High-Capacity RNA-to-cDNA kit (ThermoFisher Scientific) was used for reverse transcription to cDNA, using an input of 200 ng and a final volume of 50 µl.

#### Analysis of UBE3A-AS expression

SYBR Green qRT-PCR was used to assess the steady state levels of cynomolgus macaque *UBE3A-AS*. Total reaction volume was 10 µl, including 2 µl of cDNA, 1X PowerUp SYBR Green Master mix (ThermoFisher Scientific), and 500 nM of forward and reverse primers [Macaque *UBE3A-AS* (Forward:5’- CCTGTGAACTTTCAACCAGGA; Reverse:5’- GGATCAGACTCCAGGCCTTC)]. *ARFGAP2* was used as an internal control [Macaque *ARFGAP2* (Forward:5’- GCGTCCATCTGAGCTTCATC; Reverse:5’- CATCATTGGCTGTGCATCCA)]. Assays were evaluated with a 5-point serial dilution curve of input to estimate efficiency, R2, and slope. Primers with efficiencies that did not fall between 75% - 125% were discarded. Cycling conditions were 2 min at 50°C, 2 min at 95°C, and 40 cycles of 15 sec at 95°C and 1 min at 60°C, with readings taken at the 60°C step of every cycle. Reactions were run on a BIO-RAD CFX96 Touch Real-Time PCR Detection System (Bio-Rad). Data was retrieved and analyzed with the BIORAD CFX Maestro software (Bio-Rad). RT-PCR reactions were analyzed as described above.

#### UBE3A SNP genotyping in cynomolgus macaques

PCR and Sanger sequencing were used to identify the T and G alleles of the cynomolgus macaque *UBE3A* gene. The total reaction volume was 25 µl, including 1 µl of genomic DNA, 500 nM of forward and reverse primers [Macaque *UBE3A* SNP (Forward:5’- AGCCAGACCCAGTACTATGC; Reverse: 5’-TCTACCCGATGCCACCAAAT)], and 12.5 µl of 2X Phire Hot Start II PCR master mix (ThermoFisher Scientific). Cycling conditions were 30 sec at 98°C, and 35 cycles of 5 sec at 98°C, 5 sec at 60°C, and 10 sec at 72°C, with a final extension of 1 min at 72°C. PCR reactions were resolved in an agarose gel and the amplicon extracted using the PureLink Quick Gel Extraction and PCR purification Combo Kit (ThermoFisher Scientific). Amplicons were sequenced at the LGT Core at Texas A&M using the same primers as for amplification.

A custom TaqMan™ assay (ANYMR2V) was used to retrospectively genotype the T and G alleles of the *UBE3A* gene in the animals. Genomic DNA was isolated from peripheral blood using the DNeasy kit (Qiagen). The TaqMan Genotyping assay was performed in a total reaction volume was 10 µl, including 1 µl of genomic DNA, 1X Gene Expression Master mix (ThermoFisher Scientific), and 1X TaqMan primer assay (ThermoFisher Scientific). Cycling conditions were 2 min at 50°C, 10 min at 95°C, and 40 cycles of 15 sec at 95°C and 1 min at 60°C, with readings taken at the 60°C step of every cycle. Reactions were run on a BIO-RAD CFX96 Touch Real-Time PCR Detection System (Bio-Rad). Data was retrieved and haplotypes (homozygous or heterozygous SNP) determined with the BIORAD CFX Maestro software (Bio-Rad). Genotypes were also visually confirmed using the raw data.

#### Digital droplet qRT-PCR analysis of cynomolgus macaque UBE3A allelic expression

The allelic ratios of *UBE3A* in cynomolgus macaque were assessed using a digital droplet qRT-PCR (ddqRT-PCR). A custom TaqMan™ quantitative RT-PCR assay was developed to amplify T and G alleles of *UBE3A* (ANYMR2V). The reaction and analysis of data were performed as described above. The allelic expression was calculated as described above.

### Statistical Analyses

Linear mixed-effect regression models were used as noted with additional covariates included in the model as fixed (e.g., to examine the effect of the independent variable and the interaction between independent variables) or random effects (e.g., to account for a repeated measure of a sample or individual). A Tukey HSD multiple comparison test was used for pair-wise comparisons between samples; a Dunnett’s multiple comparison was used for multiple comparisons relative to control; indicator function parameterization test was used to compare slopes between groups; Goodness-of-fit of models were assessed by visual inspection of diagnostic residual plots, the coefficient of determination (R^2^), and AIC and BIC values. Multiplicity of testing was used for post-hoc analyses as noted. The half maximal inhibitory concentration [IC_50_ (*UBE3A-AS*)] was determined by fitting nonlinear curves with the log molar concentration of treatment as the predictor formula and ΔΔCq values as the response variable with treatment as a grouping variable to compare models across treatments. Three- or four-parameter-logistic regression models (Hill) were then fitted to determine the IC_50_. To compare treatments, a Parallelism test (F and Chi-Square tests) was run to determine whether curves were similar in shape. For similar curves, the parallel fit parameter estimates and relative potencies across treatments were reported. Statistical analyses were performed in JMP Pro 15.

## Acknowledgments

We are grateful to Ryan Doan, Bradley Revell, Cristina Cardona Barreña, and Niamh Nichols for assisting with the project and Janie Hurley for her guidance and advice. The authors would also like to thank Texas A&M Institute for Genome Sciences and Society (TIGSS) for providing computational resources and systems administration support for the TIGSS High-Performance Computing Cluster (TIGSS-HPC).

## Funding

This work was supported by the Foundation for Angelman Syndrome Therapeutics (FAST) and GeneTx Biotherapeutics, LLC.

## Author contributions

SVD was involved in Conceptualization, Data curation, Formal Analysis, Funding acquisition, Investigation, Methodology, Project administration, Resources, Supervision, Validation, Visualization, Writing – original draft, and Writing – review & editing. SGBC was involved in Formal Analysis, Funding acquisition, Investigation, Methodology, Project administration, Supervision, Validation, Visualization, and Writing – review & editing. WJM was involved in Formal Analysis, Writing – original draft, and Writing – review & editing. AB and JP were involved in Conceptualization, Data curation, Methodology, Project administration, and Resources. AS was involved in Investigation and Writing – review & editing. JB, KR, RR, LM, TJ, HS, PH, AH, and KB were involved in Investigation. KK provided Formal analysis and Investigation. LB and JD were involved in Conceptualization, Data curation, Methodology, and Resources. Members of the FIRE consortium were involved in Supervision.

## Competing interests

SVD is an employee of Ultragenyx and has an equity interest in the company. SVD, AB, and JP have an equity interest in GeneTx Biotherapeutics, LLC. SVD has a patent-pending related to this work.

## Data and materials availability

All data associated with this study are either present in the manuscript, supplementary materials, public database, or available upon request.

## #FIRE Consortium Members

Edwin Weeber - Department of Molecular Pharmacology and Physiology, University of South Florida, Tampa, FL, USA

David Segal - MIND Institute, Genome Center, and Department of Biochemistry and Molecular Medicine, University of California Davis, Davis, CA, USA

Anne Anderson - Department of Pediatrics and Neurology, Baylor College of Medicine, Houston, TX, USA

Kevin Nash - Department of Molecular Pharmacology and Physiology, University of South Florida, Tampa, FL, USA

Jill Silverman - MIND Institute and Department of Psychiatry and Behavioral Sciences, University of California Davis School of Medicine, Sacramento, CA, USA

## Notes

### Competing Interest Statement

SVD is an employee of Ultragenyx Pharmaceutical and has an equity interest in the company.  SVD, AB, and JP have an equity interest in GeneTx Biotherapeutics, LLC.  SVD has a patent-pending related to this work.

