## Supplementary Materials for "Development of an ASO therapy for Angelman syndrome by targeting an evolutionarily conserved region at the start of the *UBE3A-AS* transcript"

### SUPPLEMENTARY FIGURES

#### Supplementary Figure 1

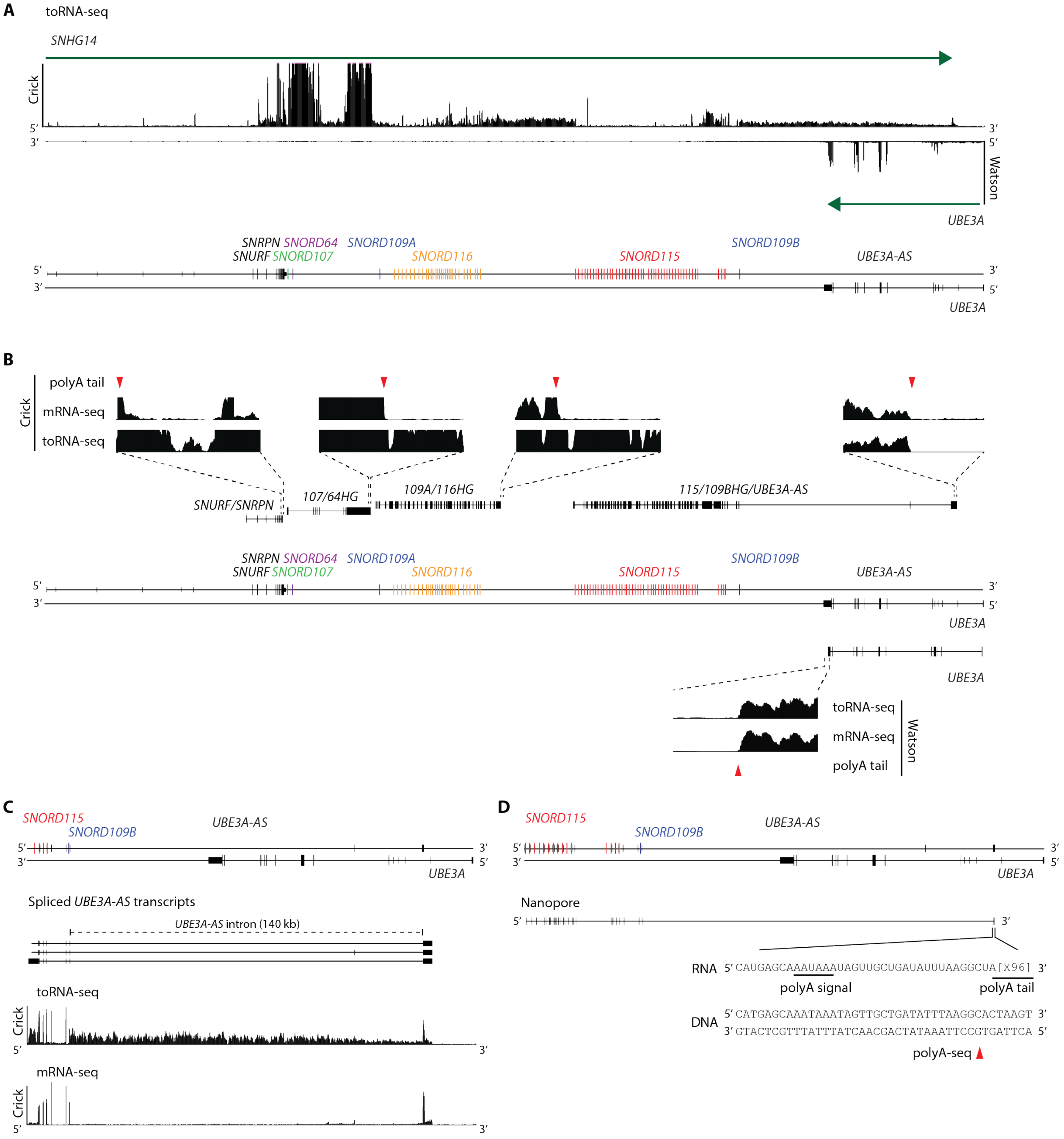

**Supplementary Fig. 1. Regulation of the *SNHG14* transcription unit.** (**A**) Inspection of the toRNA-seq reads aligned to the Crick (*SNHG14*) and Watson (*UBE3A*) strands indicates that *SNHG14* is transcribed as a single, long transcript that extends from the 5'-end of the *SNURF*/*SNRPN* to the 5'-end of *UBE3A*. (**B**) Inspection of the aligned toRNA-seq, mRNA-seq, and polyA-seq data sets in conjunction with the mRNA-seq assembled transcripts indicates that the *SNHG14* nascent transcript is alternatively spliced and processed into separate transcript units. The mRNA-seq aligned reads are depleted downstream of the polyA tails (red triangles), whereas the toRNA-seq aligned reads are present across the entire *SNHG14* transcription unit, indicating the nascent transcript is co-transcriptionally processed by 3' cleavage and polyadenylation. The black boxes and lines represent exons and introns, respectively. (**C**) The mRNA-seq assembled transcripts show the major splice variant of the *UBE3A-AS* transcript has a large intron that spans from the distal host gene exon of *SNORD109B* to the 3'-terminal exon at the 5'-end of *UBE3A*. Comparisons between the toRNA-seq and mRNA-seq aligned reads indicate the host gene and *UBE3A-AS* transcript are co-transcriptionally spliced. The toRNA-seq reads aligned to the large *UBE3A-AS* intron slope downward (5'-3') and are disproportionately higher than the smaller introns upstream. The toRNA-seq and mRNA-seq read-depth coverages are similar at the 3'-terminal exon of the *UBE3A-AS* transcript, indicating that most of the transcripts are polyadenylated. (**D**) Long-read Oxford Nanopore sequencing verifies that the *115*/*109BHG*/*UBE3A-AS* transcript is spliced and polyadenylated. The 3'-terminal exon is cleaved and polyadenylated downstream of a canonical polyA-signal (AAUAAA). The sequenced polyA-tail corresponds to a polyadenylation site in the polyA-seq data (red triangle).

#### Supplementary Figure 2

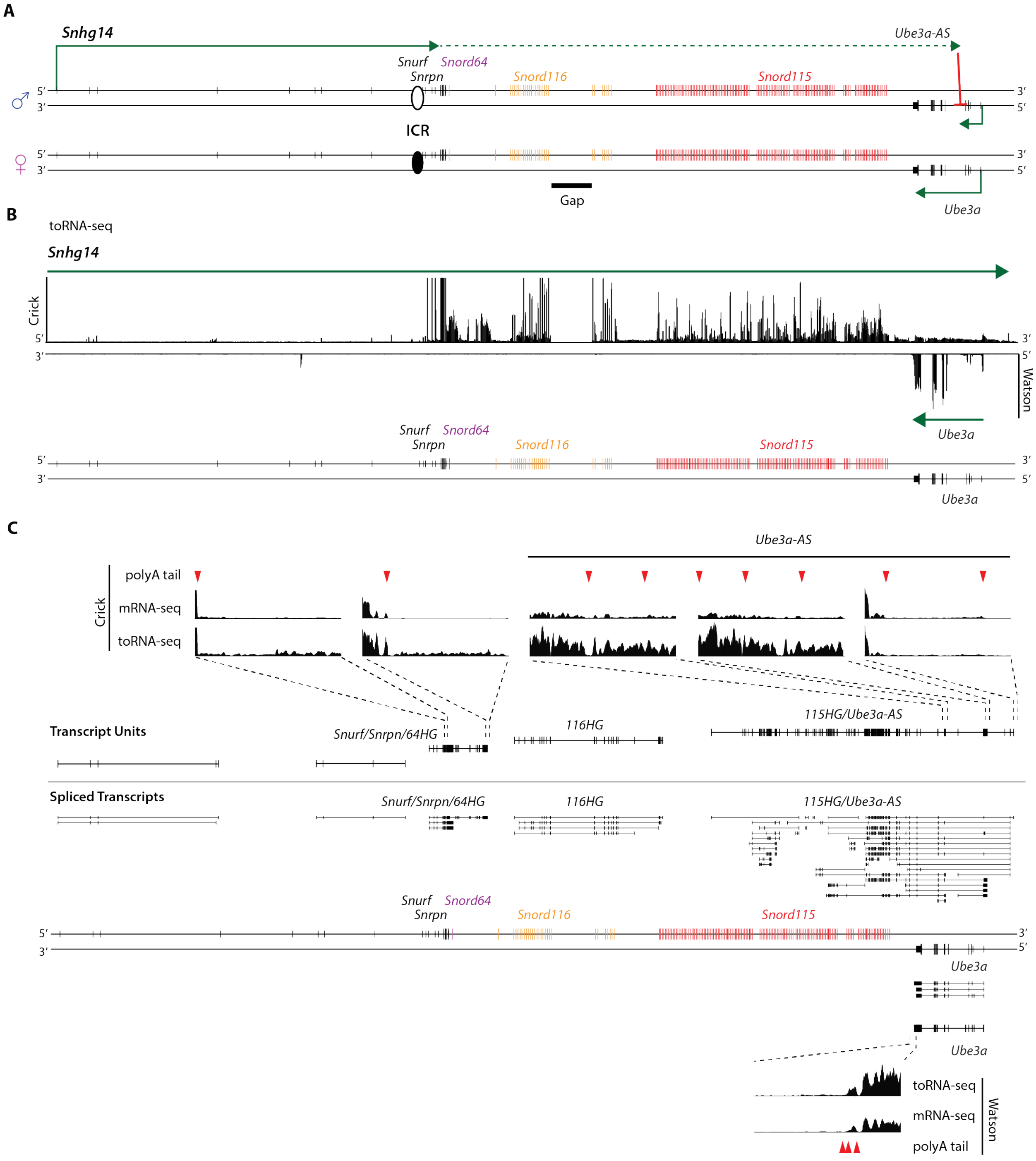

**Supplementary Fig. 2. The mouse *Ube3a-AS* transcript is a spliced, polyadenylated transcript regulated as part of the host gene.** (**A**) Schematic illustrating the imprinted and tissue-specific regulation of the mouse *Snhg14* transcription unit and *Ube3a*. The *Snhg14* polycistronic transcript is comprised of the *Snurf/Snrpn* protein-coding mRNAs, *Snord107*, *Snord64*, *Snord116* (71 copies), *Snord115* (136 copies), and the *Ube3a-AS* transcript. The *Snord116* gene array is not fully assembled in the mouse genome (Gap). (**B**) Inspection of the toRNA-seq reads aligned to the Crick (*Snhg14*) and Watson (*Ube3a*) strands indicates that *Snhg14*is transcribed as a single, long transcript that extends from the 5'-end of the *Snurf/Snrpn* to a region upstream of *Ube3a*. The proximal end of *Snhg14* is transcribed in all tissues (solid line), whereas the distal end is only transcribed in CNS neurons (dashed line), which represses the paternal *Ube3a* allele. (**C**) Inspection of the aligned toRNA-seq, mRNA-seq, and polyA-seq data sets in conjunction with the mRNA-seq assembled transcripts indicates that the *Snhg14* nascent transcript is processed into separate transcript units by 3' cleavage and polyadenylation. The mRNA-seq aligned reads are reduced or depleted downstream of the polyA tails (red triangles), whereas the toRNA-seq aligned reads are abundant and present across the entire *Snhg14* transcription unit. The black boxes and lines represent exons and introns, respectively.

#### Supplementary Figure 3

**A**

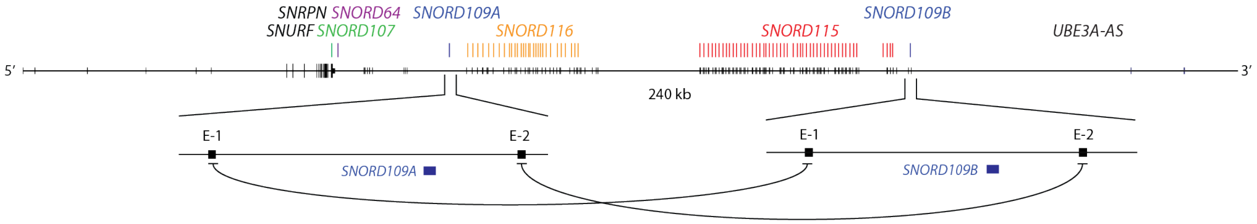


**B**

*SNORD109 E-1*

| | | size (nt) | Percent Identity |
|---|---|---|---|
| *SNORD109B* | AGGT-TAAAAGCTGAAACAACTTCAAGGAAACTTCAGGGAAAAGAGGT | 47 | |
| *SNORD109A* | AGGTCTAGAAGATGTCACAG---------AATTTCAAGGAAGAGAGGT | 39 | 62.5 |

*SNORD109 E-2*

| | | size (nt) | Percent Identity |
|---|---|---|---|
| *SNORD109B* | AGA--AGGCCTGGAATCTGATCCTCCACTTCAGAGAACAGGGGT | 42 | |
| *SNORD109A* | AGAGGAGGGCTGGAATGAGGTCCTCCACTTCAGAAAATAGGGGT | 44 | 81.82 |

**C**

*SNORD109*

| | | size (nt) | Percent Identity |
|---|---|---|---|
| *SNORD109B* | GGATCGATGATGAGAATAATTGTCTGAGGATGCTGAGGGACTCATTCCAGATGTCAATCTGAGGTCC | 67 | |
| *SNORD109A* | GGATCGATGATGAGAATAATTGTCTGAGGATGCTGAGGGACTCATTCCAGATGTCAATCTGAGGTCC | 67 | 100% |

**Supplementary Fig. 3. The *SNORD109A* and *SNORD109B* genes and host gene exons are duplicated at the distal and proximal ends of the *SNORD116* and *SNORD115* gene arrays.** (**A**) Schematic illustrating duplication of *SNORD109A* and *SNORD109B* genes and host gene exons (E-1 and E-2) at the proximal and distal ends of the *SNORD116* and *SNORD115* gene arrays. The annotated *SNORD109B* host exons are homologous to unannotated sequences flanking *SNORD109A*. (**B and C**) Sequence alignments of the *SNORD109A/B* host exons and gene sequences.

#### Supplementary Figure 4

**A**

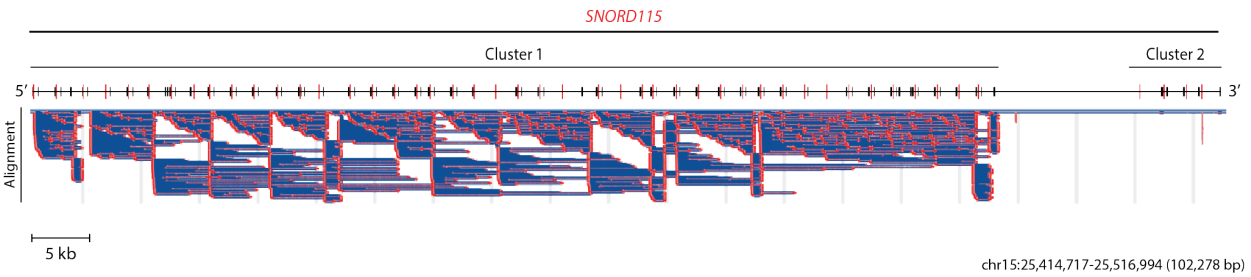

**Supplementary Fig. 4. The *SNORD115* gene array is partitioned into two clusters. (A)** Schematic showing repeated units in Cluster 1. A discontinuous megablast self-alignment of the *SNORD115* gene array shows that Cluster 1 is organized as repeated units ranging from 1.9 to 5 kb in size with varying degrees of sequence homology. In contrast, Cluster 2 exists as a single copy with no repeated structures, apart from the *SNORD115-48* gene, which is homologous to the *SNORD115* genes in Cluster 1. The red and blue alignments indicate highly and moderately conserved sequences, respectively.

### Supplementary Figure 5

**A**

**Cluster 1**

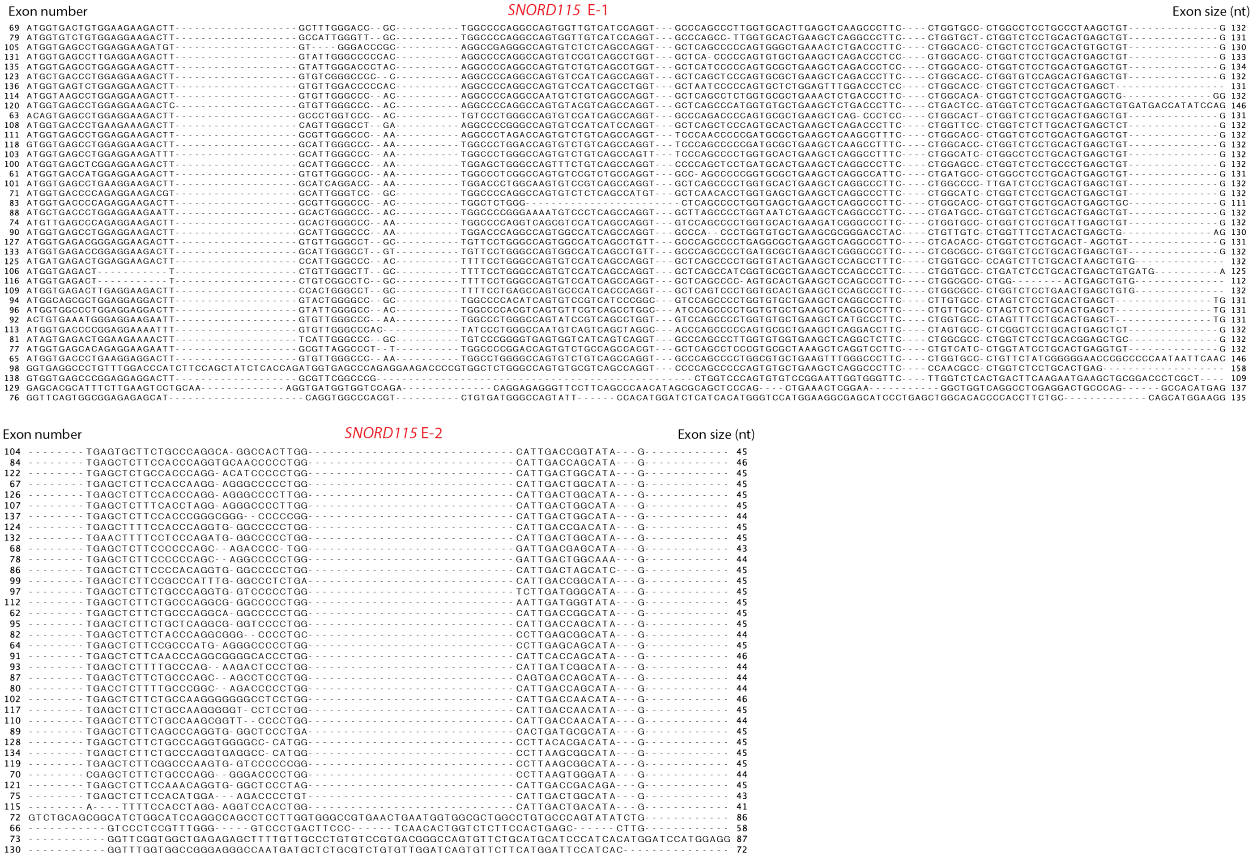


**B**

**Cluster 2**

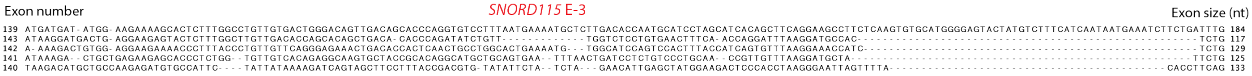

**Supplementary Fig. 5. The host gene exons in Clusters 1 and 2 are divergent. (A - B)** Sequence alignments of the E-1 and E-2 host gene exons in Cluster 1 and the E-3 host gene exons in Cluster 2.

### Supplementary Figure 6

**A**

*SNORD115*

```
                    Stem     C Box                                                     Antisense element        D Box    Stem   size (nt)
     1 GTGTTGATGATGAGAACCTTATATTAT-CCTGAAGAGAGGTGATGACTT-AAAA--ATCATGCTCAATA---GGATTACGCTGAGGCCC 82
     2 GGGTCGATGATGAGAAGCTTCTGTTTT-CTTGAAGAGAGGTGATGACTT-AAAA--ATCATGCTCAATA---GGATTATGCTGAGGCCC 82
     3 GGGTCAATGATGAGAACCTTATATTGT-CCTGAAGAGAGGTGATGACTT-AAAA--ATCATGCTTAGTA---GGATTACGCTGAGGCCT 82
     4 GGGTCGATGATGAGAACTTTATATTGT-TCTGAAGAGAGGTGATGACTT-AAAA--ATCATGCTCAATA---GGATTACGCTGAGGCCC 82
     5 GGATCGATGATGAGAACCTTATATTGT-CCTGAAGAGAGGTGATGACTT-AAAA--ATCATGCTCAATA---GGATTACGCTGAGGCCC 82
     6 GGGTCAATGATGAGAACCTTATATTGT-TCTGAAGAGAGGTGATGACTT-AAAA--ATCATGCTCAATA---GGATTACGCTGAGGCCC 82
   * 7 GGGTCAATGA---GAACCTTATATTGT-CCTGAAGAGAGGTGATAACTT-AAAA--ATCATGCTCAATAATAGGATTACGCTGAGGCCC 82
     8 GGGTCAATGATGAGAACCTTACATTGT-TCTGAAGAGAGATGATGACTT-AAAA--ATCATGCTCAATA---GGATTACGCTGAGGCCC 82
     9 GGGTCGATGATGAGAACCTTATATTGT-CCTGAAGAGAGGTGATGACTT-AAAA--ATCATGCTCAATA---GGATTACGCTGAGGCCC 82
    10 GGGTCGATGATGAGAACCTTATATTGT-C-TGAAGAGAGGTGATGACTT-AAAA--ATCATGCTCAATA---GGATTACGCTGAGGCCC 81
    11 GGGTCAATGATGAGAACCTTATATTGT-CCTGAAGAGAGGTGATGACTT-AAAA--ATCATGCTCAATA---GGATTACGCTGAGGCCC 82
    12 GGGTCGATGATGAGAACCTTATATTGT-CCTGAAGAGAGGTGATGACTT-AAAA--ATCATGCTCAATA---GGATTACGCTGAGGCCC 82
    13 GGGTCGATGATGAGAACCTTATATTAT-CCTGAAGAGAGGTGATGACTT-AAAA--ATCATGCTCAATA---GGATTACGCTGAGGCCC 82
    14 GGGTCGATGATGAGAAACTTATATTGT--CTGAAGAGAGGTGATGACTT-AAAA--ATCATGCTCAATA---GGATTACGCTGAGGCCC 81
    15 GGGTCGATGATGAGAACCTTATAT-GT-TCTGAAGAGAGGTGATGACTT-AAAA--ATCATGCTCAATA---GGATTACGCTGAGGCCC 81
    16 GGGTCAATGATGAGAACCTTATATTAT-CCTGAAGAGAGGTGATGACTT-AAAA--ATCATGCTCAATA---GGATTACGCTGAGGCCC 82
    17 GGGTCGATGATGAGAACCTTATATTGT-CCTGAAGAGAGGTGATGACTT-AAAA--ATCATTCTCAAAA---GGATTATGCTGAGGCCC 82
    18 GGGTCGATGATGAGAACCTTATATTGT-CCTGAAGAGAGGTGATGACTT-AAAA--ATCATTCTCAAAA---GGATTATGCTGAGGCCC 82
    19 GGGTCGATGATGAGAACCTTATATTGT-CCTGAAGAGAGGTGATGACTT-AAAA--ATCATTCTCAAAA---GGATTATGCTGAGGCCC 82
    20 GGGTCGATGATGAGAACCTTATATTGT-CCTGAAGAGAGGTGATGACTT-AAAA--ATCATGCTCAATA---GGATTATGCTGAGGCCC 82
    21 GGGTCGATGATGAGAACCTTATATTTT-C-TGAAGAGAGGTGATGACTT-AAAA--ATCATGCTCAATA---GGATTACGCTGAGGCCC 81
    22 GGGTCAATGATGAGAACCTTATATTGT-CCTGAAGAGAGGTGATGACTT-AAAA--ATCATGCTCAATA---GGATTACGCTGAGTCCC 82
    23 GGGTCAATGATGAGAACCCTATATTGT-GTTGAAGAGAGGTGATGACTT-AAAATTACCATGCTCAAT-----GATTACGCTGAGGCCC 82
  * 24 G-----------AGAACCTTATATTGT-TCTGAAGAGAGGTGGTGACTT-AAAA--ATCATGCTCAATA---GGATTACGCTGAGGCCC 71
  * 25 AGGTCGATTATGAGAACCTTATATTGT-CCTGAAGAGAGGTGATGACTT-AAAA--ATCATGCCCAATA---GGATTACGCTGAGGCCC 82
    26 GGGTCAGTGATGAGAACCTTATATTGT-CCTGAAGAGAGGTGATGACTT-AAAA--ATCATGCTCAATA---GGATTACGCTGAGGCCC 82
  * 27 ------ATGATGAGAACCTTATATTGT-CCTGAAAAGAGGTGATGACTT-AACA--ATCATGCTCAATA---GGATTACATTGAAGCCC 76
  * 28 -------TGATGAGAACCTTGTATTCT-TCTGAAGAGAGGTGATGACTT-AAAA--ACCATGCTCAATA---GGATTACACTTAGGCC- 74
    29 GGGTCAATGATGAGAACCTTATATTGT-CCTGAAGAGAGGTGATGACTT-AAAA--ATCATGCTCAATA---GGATTACGCTGAGGCCC 82
    30 GGGTCAATGATGAGAACCTTATATTGT-TCTGAAGAGAGGTGATTATTT-AAAA--ATCATGCTCAATA---GGATTACGCTGAGGCCC 82
    31 GGGTCAGTGATGAGAACCTTATATTGT-CCTGAAGAAAGGTGATGACTT-AAAA--ATCATGCTCAATA---GGATTACACTGAGGCCC 82
    32 GGGTCAATGATGAGAACCTGATATTGC-CCTGAAGAGAGATGATGACTT-AAAA--ATCATGTTCAATA---GGATTACGCTGAGGCCT 82
    33 GGGTCAATGATGAGAACCGTATATTGT-CCTGAAGAGCGGTGATGACTT-AAAA--ATAATGCTCAATA---GGATTACGCTGAGGCCC 82
    34 GGGTCAATGATGAGAACCTTATAATGT-TCTGAAGAGAGGTGATGACTT-AAAA--ATCATGCTCAATA---GGATTACGCTGAGGCCC 82
    35 GGGTCAATGATGAGAACCTTGTATTAT-CTTGAAGAGAGGTGATGACTT-AAAA--ATCATGCTCAATA---GGATTACACTGAGGCCC 82
    36 GGGTCAATGATGAGAACCTTATATTGT-CCTGAAGAGAGGTGATGACTT-AAAA--ATCATGCTCAATA---GGATTACGCTGAGGCCC 82
    37 GGGCTGATGATGAGAACCTTATATTGT-CCTGAAAAAAGGTGATGACTT-AAA--CATCATGCTTAATA---GTATTATGCTGAAGCCC 82
    38 GGGTCAATGATGAGAACCTTACATTGT-CCTGAAGAGAGATGATGACTT-AAAA--ATCATGCTCAATA---GGATTACGCTGAGGCCC 82
    39 GGGTCAATGATGAGAATCTTATATTGT-CCTGAAGAGAGGTGATGACTT-AAAA--ATCATGCTCAATA---GGATTACGCTGAGGCCC 82
    40 GGGTCGATGATGAGAACCTTATATTTT-CCTGAAGAGAGGTGATGACTT-AAAA--ATCATGCTCAATA---GGATTACGCTGAGGCCC 82
    41 GGGTCAGTGATGAGAACCTTCTATTGT-CCTGAAGAGAGGTGATGACTT-AAAA--ATCATGCTCAATA---GGATTACGCTGAGGCCC 82
    42 GGGTCGATGATGAGAACCTTATATTGT-TCTGAAGAGAGGTGATGACTT-AAAA--ATCATGCTCAATA---GGATTACGCTGAGGCCC 82
    43 GGGTCAATGATGAGAACCTTATATTGT-CCTGAAGAGAGGTGATGACTT-AAAA--ATCATGCTCAATA---GGATTACGCTGAGGCCC 82
    44 GGGTCAATGATGAGAACCTTATATTGT-CCTGAAGAGCGGTGATGACTT-AAAA--ATCATGCTCAATA---GGATTACGCTGAGGCCC 82
  * 45 GGGGCAAT-ATGGAGATGTTATATTGT-CTTCGACAGGGAAGATGACAT-AAAA--ATTATGTTCAATA---GGATTATGTGGAGACTT 81
 *45a ------ATGACAAAATCTTTACTTTTTATCTGAA------------------------------------------------------- 28
  * 46 ----CAATAATGAAATCTTC----TGA-TTTGGTGAGAAATAATGCCTT-AAAA---TTACACTCAATA---GGATTATGCTGAGGC-- 71
  * 47 -----AAATATATGTATAA-AGAGTAT-CTTCAGGAGACGTAATAATGT-AAAA--ATCATGCTCAATA---GAATTAAGCTGAGGCTC 76
    48 GGGTCAATGATGAGA------TGTTAC-CTTGAAGAGAAATGATGACGT-AAAA--ATTAAGTTCAGT---TGGATTACGCTGAGGCCC 76
  * 49 GTTTCAACAATAAGA-CCTTATGTTGT-CTTGAGGACATGAGATGACATGAAAACTAGACTGGCTGGTCCTTATCTTTT-CTGGGG--- 83
```

Cluster 1: 1–44

Cluster 2: 45–49

**Supplementary Fig. 6. The *SNORD115* genes in Clusters 1 and 2 are divergent.** (**A**) Sequence alignment of *SNORD115* genes in Cluster 1 and Cluster 2. The conserved C/D box motifs are underlined. The *SNORD115-45a* and *SNORD115-49* pseudogenes were identified by sequence analysis. *SNORD115* genes with degenerate C/D box motifs (pseudogenes) are indicated by an asterisk (*).

### Supplementary Figure 7

**A**

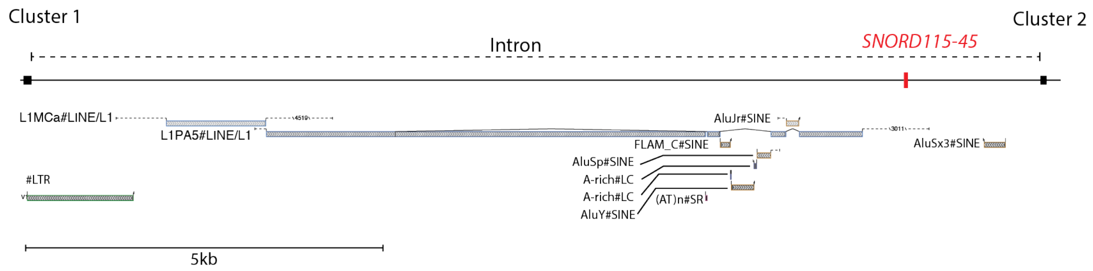

**Supplementary Fig. 7. Cluster 1 is separated from Cluster 2 by an atypical host gene intron harboring an array of repetitive elements.** (**A**) Schematic of repetitive elements in the intron separating Clusters 1 and 2.

### Supplementary Figure 8

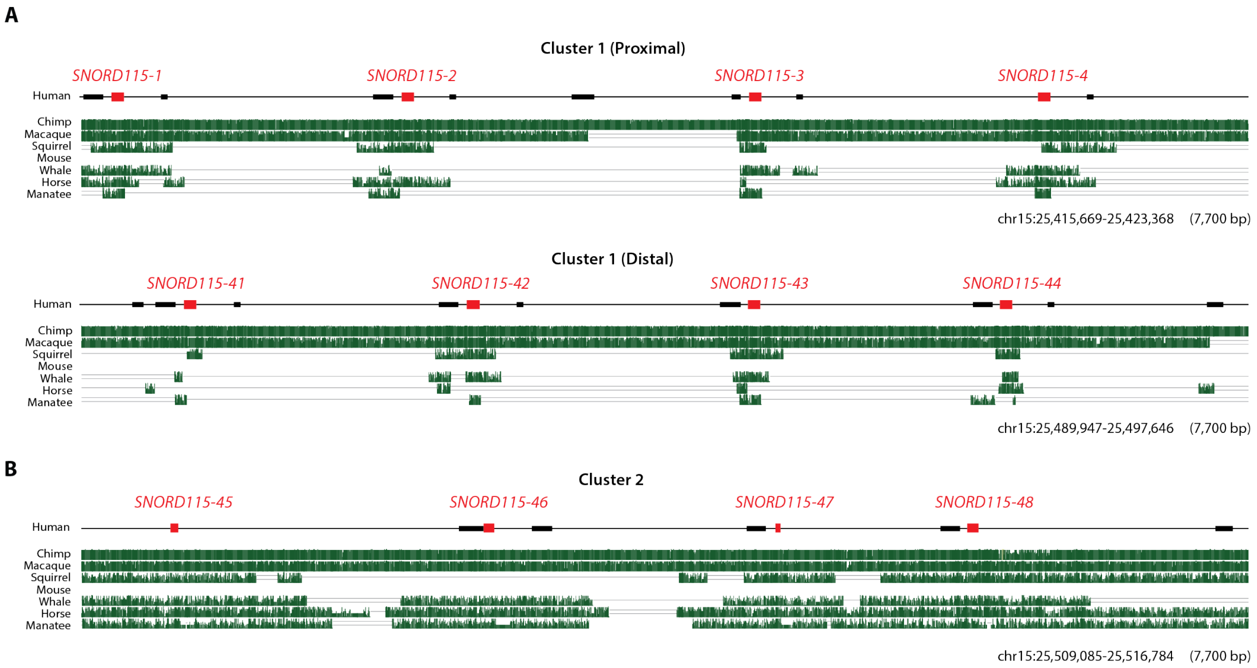

**Supplementary Fig. 8. Clusters 1 and 2 have been subjected to different evolutionary constraints.** (**A**) Multiz genome alignments between human and the orthologous regions in distantly related animals at the proximal and distal ends of Cluster 1 show short stretches of conserved sequences (vertical green lines) corresponding to the *SNORD115* gene sequences. (**B**) Multiz genome alignments between human and the orthologous regions in distantly related animals in Cluster 2 show large blocks of conserved sequences spanning the region, except for the mouse genome, which has uniquely lost the region.

#### Supplementary Figure 9

**A**

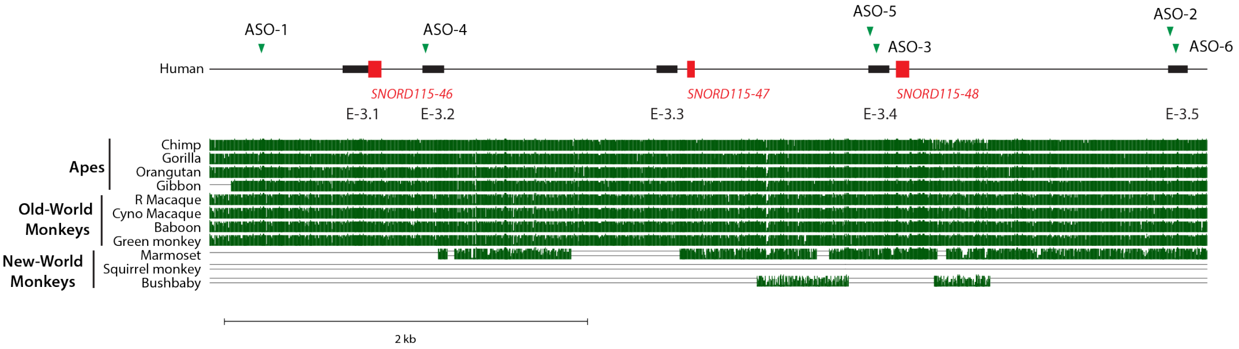


**B**

| | | Conserved ASOs | | |
|---|---|---|---|---|
| | | **ASO-3** | **ASO-4** | **ASO-6** |
| | Human | TGTTCAGGGAGAAACTGACA | CCAAGAGATGTGCCATTCTA | CTTTGGCTTGTTGACACCAG |
| **Apes** | Chimp | TGTTCAGGGAGAAACTGACA | CCAAGAGATGTGCCATTCTA | CTTTGGCTTGTTGACACCAG |
| | Bonobo | TGTTCAGGGAGAAACTGACA | CCAAGAGATGTGCCATTCTA | CTTTGGCTTGTTGACACCAG |
| | Gorilla | TGTTCAGGGAGAAACTGACA | CCAAGAGATGTGCCATTCTA | CTTTGGCTTGTTGACACCAG |
| | Orangutan | TGTTCAGGGAGAAACTGACA | CCAAGAGATGTGCCATTCTG | CTTTGGCTTGTTGACACCAG |
| | Gibbon | TGTTCAGGGAGAAACTGACA | CCAAGAAATGTGCCATTCTG | CTTTGGCTTGTTGACACCAG |
| **Old-World Monkeys** | Rhesus | TGTTCAGGGAGAAACTGACA | CCAAGAGATGTGCCATTCTA | CTTTGGCTTGTTGACACCAG |
| | Cyno Macaque | TGTTCAGGGAGAAACTGACA | CCAAGAGATGTGCCATTCTA | CTTTGGCTTGTTGACACCAG |
| | Pig-tailed Macaque | TGTTCAGGGAGAAACTGACA | CCAAGAGATGTGCCATTCTA | CTTTGGCTTGTTGACACCAG |
| | Sooty Mangabey | TGTTCAGGGAGAAACTGACA | CCAAGAGATGTGCCATTCTA | CTTTGGCTTGTTGACACCAG |
| | Baboon | TGTTCAGGGAGAAACTGACA | CCAAGAGATGTGCCATTCTG | CTTTGGCTTGTTGACACCAG |
| | Green Monkey | TGTTCAGGGAGAAACTGACA | CCAAGAGATGTGCCATTCTA | CTTTGGCTTGTTGACACCAG |
| | Drill | TGTTCAGGGAGAAACTGACA | CCAAGAGATGTGCCATTCTA | CTTTGGCTTGTTGACACCAG |
| | Proboscis Monkey | TGTTCAGGGAGAAACTGACA | CCAAGAGATGTGCCATTCTA | CTTTGGCTTGTTGACACCAG |
| | Angolan Colobus | TGTTCAGGGAGAAACTGACA | CCAAGAGATGTGCCATTCTA | CTTTGGCTTGTTGACACCAG |
| | Golden Snub-Nosed Monkey | TGTTCAGGGAGAAACTGACA | CCAAGAGATGTGCCATTCTA | CTTTGGCTTGTTGACACCAG |
| | Black Snub-Nosed Monkey | TGTTCAGGGAGAAACTGACA | CCAAGAGATGTGCCATTCTA | CTTTGGCTTGTTGACACCAG |

**Supplementary Fig. 9. Conservation of ASO target sequences across humans, apes, and Old-World monkeys.** (**A**) Multiz genome alignments between human and the orthologous regions in apes and Old-World monkeys. Conserved sequences are depicted by vertical green lines. The ASO target regions are depicted by green triangles. (**B**) Sequence alignments show that the ASO-3, ASO-4, and ASO-6 target sequences are identical across humans, apes, and Old-World monkeys, indicating these sequences have been evolutionarily constrained for at least 29 million years.

### Supplementary Figure 10

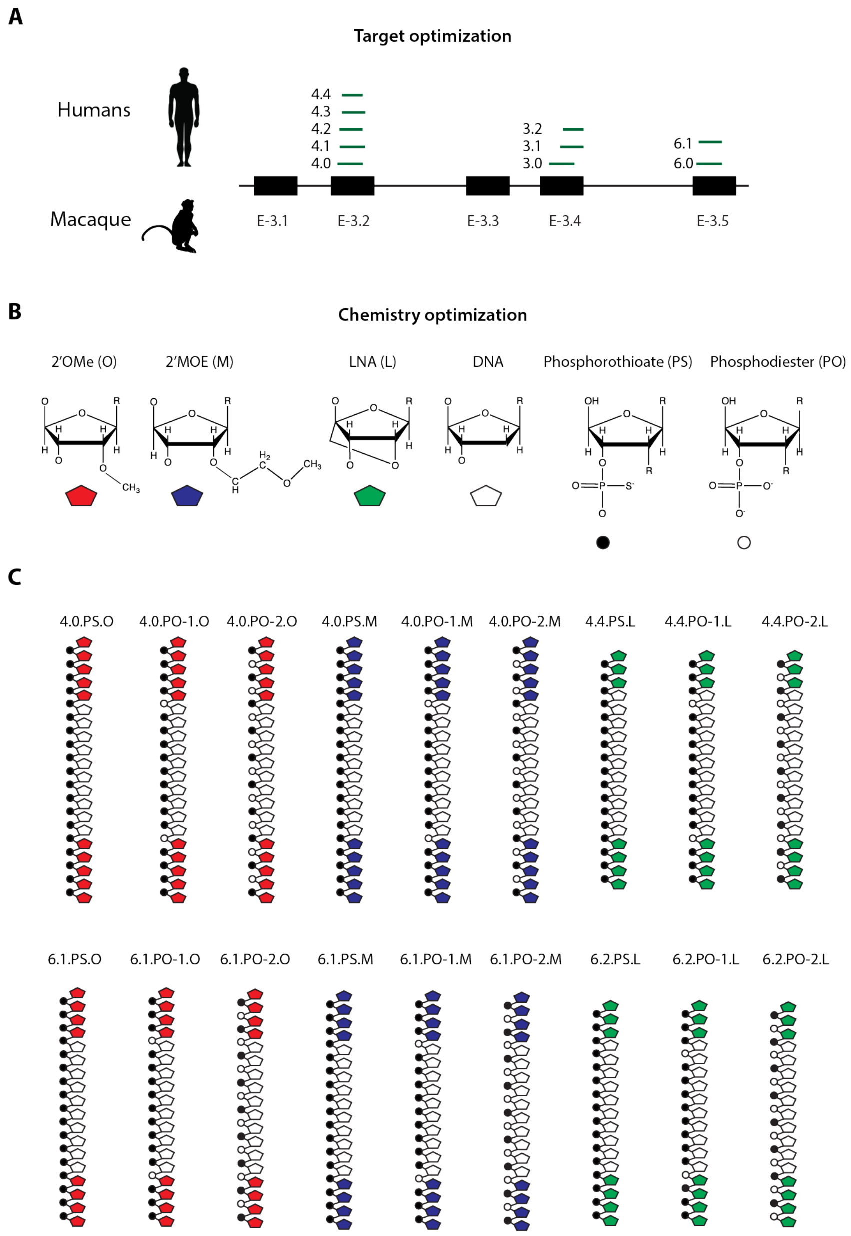

**Supplementary Fig. 10. Schematic of sequence and chemically optimized ASOs.** (**A**) Schematic illustrating the strategy used to optimize the target sequences of the conserved ASOs (ASO-3, ASO-4, and ASO-6). Only identical sequences across humans and between human and cynomolgus macaque were targeted. (**B**) Schematic of the modified ribonucleosides [2'OMe (O), 2'-MOE (M), LNA (L)], a deoxynucleoside (DNA), and phosphorothioate (PS) and phosphodiester (PO) linkages. (**C**) Schematics of the chemically optimized ASOs. The modified ribonucleosides and deoxynucleosides are depicted by colored (red, blue, and green) and white pentagons, respectively; the phosphorothioate and PO phosphodiester are depicted by black and white circles, respectively.

#### Supplementary Figure 11

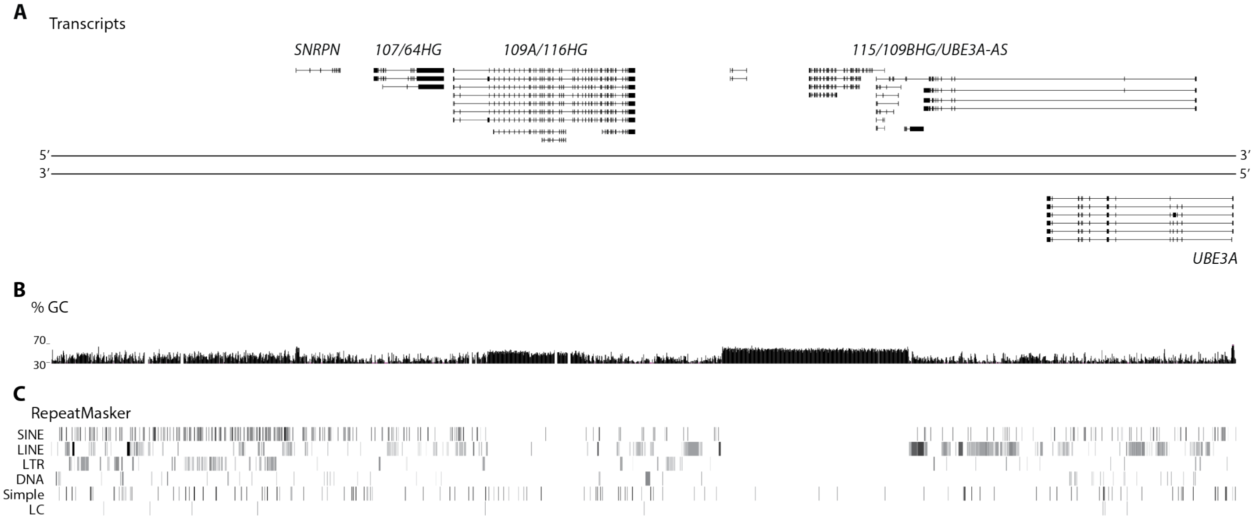


macFas5 chr7:2,805,112-3,442,080

**Supplementary Fig. 11. The cynomolgus macaque *SNHG14* transcript is processed like the human *SNHG14* transcript.** (**A**) Transcripts assembled from mRNA-seq performed on cynomolgus macaque motor cortex (n = 1). (**B and C**) Percentage of guanine and cytosine nucleotides [% GC (scale = 30 — 70%)] and distribution of repetitive elements. Abbreviations: SINE, small interspersed nuclear elements; LINE, long interspersed nuclear elements; LTR, long terminal repeat; DNA, DNA repeats; Simple, simple repeats; LC, low copy repeats.

#### Supplementary Figure 12

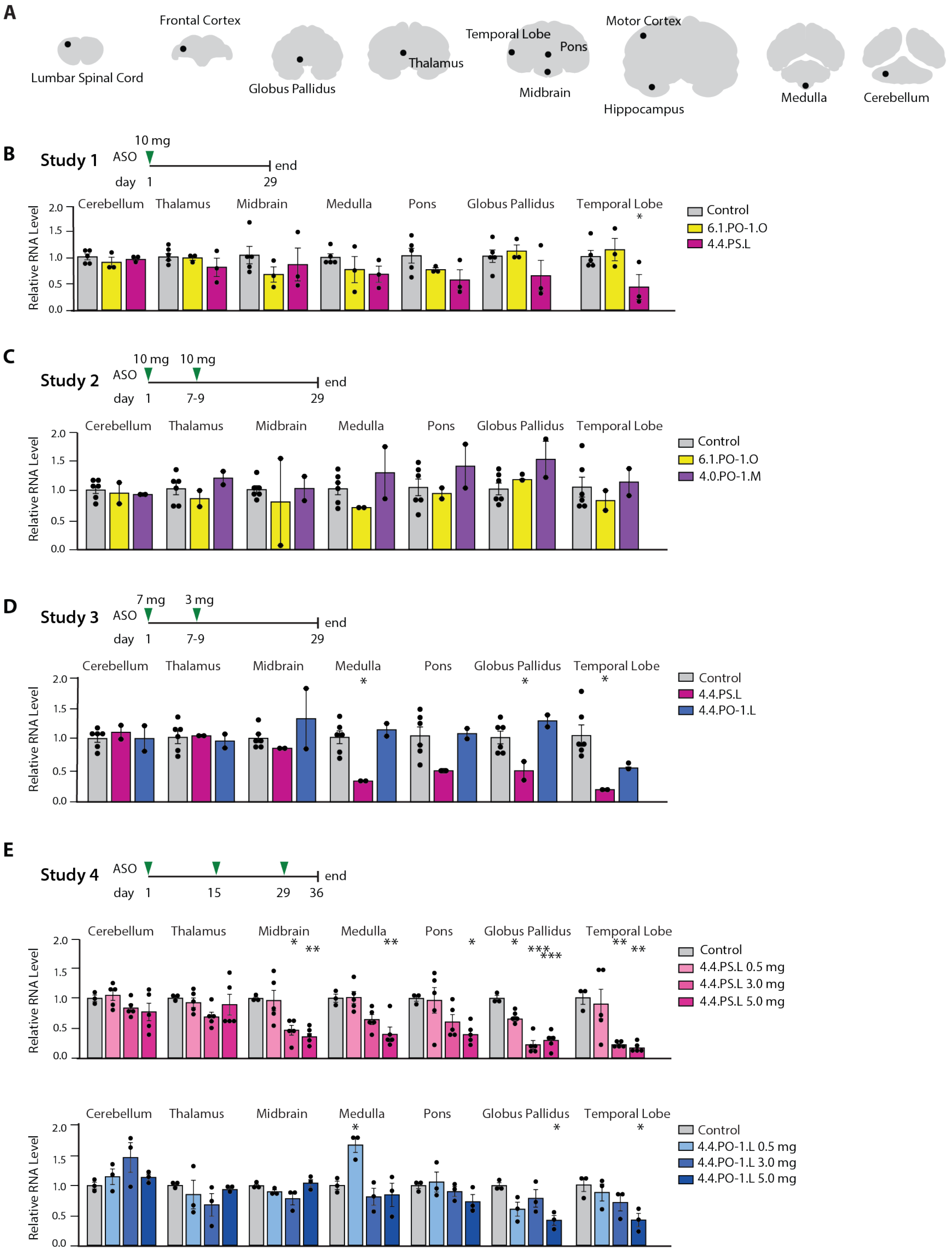

**Supplementary Figure 12. ASO treatment reduces *UBE3A-AS* RNA levels in the cynomolgus macaque CNS.** (**A**) Schematic of CNS regions analyzed in cynomolgus macaques. (**B**) *UBE3A-AS* RNA levels after a single injection of 6.1.PO-1.O (10 mg, n = 3) and 4.4.PS.L (10 mg, n = 3), normalized to control (n = 5) (dashed line). Data presented as mean ± SEM; Mixed effect linear regression followed by Dunnett's test, *$P < 0.05$, **$P < 0.01$, ***$P < 0.0001$. (**C**) Experimental timeline of ASO treatment and analysis. *UBE3A-AS* RNA levels after a two injections of 6.1.PO-1.O [10 mg, 10 mg (n = 3)] and 4.0.PO-1.M [10 mg, 10 mg (n = 3)], normalized to control (n = 6). Data presented as mean ± SEM; Mixed effect linear regression followed by Dunnett's test. *$P < 0.05$, **$P < 0.01$, ***$P < 0.0001$. (**D**) *UBE3A-AS* RNA levels after a two injections of 4.4.PS.L [7 mg, 3 mg (n = 2)] and 4.4.PO-1.L [7 mg, 3 mg (n = 2)], normalized to control (n = 6). Data presented as mean ± SEM; Mixed effect linear regression followed by Dunnett's test. *$P < 0.05$, **$P < 0.01$, ***$P < 0.0001$. (**E**) Experimental timeline of ASO treatment and analysis. *UBE3A-AS* RNA levels after three injections of 4.4.PS.L [0.5 mg, 3.0 mg, and 5.0 mg (n = 5/dose)], normalized to control (n = 3). Data presented as means ± SEM; Mixed effect linear regression followed by Dunnett's test. *$P < 0.05$, **$P < 0.01$, ***$P < 0.0001$. (**F**) *UBE3A-AS* RNA levels after three injections of 4.4.PO-1.L [0.5 mg, 3.0 mg, and 5.0 mg (n = 3/dose)], normalized to control (n = 3). Data presented as means ± SEM.; Mixed effect linear regression followed by Dunnett's test. *$P < 0.05$, **$P < 0.01$, ***$P < 0.0001$.

#### SUPPLEMENTARY TABLES

**Table S1. Human ASOs**

| ASO | RNA Modification | Backbone | Size (nt) | Design (5´-3´) | Sequence (5’-3’) |
|---|---|---|---|---|---|
| Control ASO | OMe | PS | 20 | 5-10-5 | $A^{O}*G^{O}*A^{O}*G^{O}*A^{O}*c*g*a*g*g*c*c*g*a*c*U^{O}*A^{O}*G^{O}*U^{O}*G^{O}$ |
| 1.0.PS.O | OMe | PS | 20 | 5-10-5 | $U^{O}*A^{O}*G^{O}*A^{O}*G^{O}*g*t*g*a*a*g*g*c*c*a*G^{O}*G^{O}*C^{O}*A^{O}*C^{O}$ |
| 2.0.PS.O | OMe | PS | 20 | 5-10-5 | $G^{O}*U^{O}*A^{O}*C^{O}*U^{O}*c*t*t*c*c*t*c*a*g*t*C^{O}*A^{O}*U^{O}*C^{O}*C^{O}$ |
| 3.0.PS.O[C] | OMe | PS | 20 | 5-10-5 | $U^{O}*G^{O}*U^{O}*C^{O}*A^{O}*g*t*t*t*c*t*c*c*c*t*G^{O}*A^{O}*A^{O}*C^{O}*A^{O}$ |
| 4.0.PS.O [C] | OMe | PS | 20 | 5-10-5 | $U^{O}*A^{O}*G^{O}*A^{O}*A^{O}*t*g*g*c*a*c*a*t*c*t*C^{O}*U^{O}*U^{O}*G^{O}*G^{O}$ |
| 5.0.PS.O | OMe | PS | 20 | 5-10-5 | $G^{O}*U^{O}*U^{O}*U^{O}*U^{O}*c*t*t*c*c*t*c*c*a*c*A^{O}*G^{O}*U^{O}*C^{O}*U^{O}$ |
| 6.0.PS.O [C] | OMe | PS | 20 | 5-10-5 | $C^{O}*U^{O}*G^{O}*G^{O}*U^{O}*g*t*c*a*a*c*a*a*g*c*C^{O}*A^{O}*A^{O}*A^{O}*G^{O}$ |

Capital letter, RNA; lower-case letter, DNA
Abbreviations: PS and *, phosphorothioate; O, OMe; C, conserved with cynomolgus macaque.

#### Table S2. Sequence Optimized ASOs

| ASO | RNA Modification | Backbone | Size (nt) | Design (5´-3´) | Sequence (5'-3') |
|---|---|---|---|---|---|
| 3.1.PS.O$^{C}$ | OMe | PS | 19 | 4-10-5 | G$^{O}$*U$^{O}$*U$^{O}$*G$^{O}$*a*g*t*g*g*t*g*t*c*a*G$^{O}$*U$^{O}$*U$^{O}$*U$^{O}$*C$^{O}$ |
| 3.2.PS.O$^{C}$ | OMe | PS | 18 | 4-10-4 | U$^{O}$*U$^{O}$*G$^{O}$*A$^{O}$*g*t*g*g*t*g*t*c*a*g*U$^{O}$*U$^{O}$*U$^{O}$*C$^{O}$ |
| 6.1.PS.O$^{C}$ | OMe | PS | 18 | 4-10-4 | C$^{O}$*U$^{O}$*G$^{O}$*G$^{O}$*t*g*t*c*a*a*c*a*a*g*C$^{O}$*C$^{O}$*A$^{O}$*A$^{O}$ |
| 4.1.PS.O$^{C}$ | OMe | PS | 20 | 5-10-5 | A$^{O}$*U$^{O}$*A$^{O}$*G$^{O}$*A$^{O}$*a*t*g*g*c*a*c*a*t*c*U$^{O}$*C$^{O}$*U$^{O}$*U$^{O}$*G$^{O}$ |
| 4.2.PS.O$^{C}$ | OMe | PS | 19 | 4-10-5 | A$^{O}$*G$^{O}$*A$^{O}$*A$^{O}$*t*g*g*c*a*c*a*t*c*t*C$^{O}$*U$^{O}$*U$^{O}$*G$^{O}$*G$^{O}$ |
| 4.3.PS.O$^{C}$ | OMe | PS | 19 | 4-10-5 | U$^{O}$*A$^{O}$*G$^{O}$*A$^{O}$*a*t*g*g*c*a*c*a*t*c*U$^{O}$*C$^{O}$*U$^{O}$*U$^{O}$*G$^{O}$ |
| 4.4.PS.O$^{C}$ | OMe | PS | 18 | 4-10-4 | A$^{O}$*G$^{O}$*A$^{O}$*A$^{O}$*t*g*g*c*a*c*a*t*c*t*C$^{O}$*U$^{O}$*U$^{O}$*G$^{O}$ |

Capital letter, RNA; lower-case letter, DNA
Abbreviations: nt, nucleotide; PS and *, phosphorothioate; O, OMe; C, conserved with macaque

**Table S3. Pharmacological Properties of Sequence Optimized ASOs**

| ASO | $IC_{50}$ (nM) | SEM | Relative Potency | $E_{max}$ (3 μM) | SEM |
|---|---|---|---|---|---|
| 4.0.PS.O | 1,300.35 | 0.00 | 1.00 | 0.31 | 0.01 |
| 4.1.PS.O | 1,985.74 | 0.05 | 0.65 | 0.42 | 0.01 |
| 4.2.PS.O | 1,091.86 | 0.08 | 1.19 | 0.31 | 0.01 |
| 4.3.PS.O | 1,858.07 | 0.05 | 0.70 | 0.40 | 0.01 |
| 4.4.PS.O | 1,883.32 | 0.05 | 0.69 | 0.41 | 0.01 |
| 3.1.PS.O | 2,991.38 | 0.03 | 0.43 | 0.51 | 0.01 |
| 3.2.PS.O | 1,839.91 | 0.05 | 0.71 | 0.44 | 0.02 |
| 6.1.PS.O | 535.09 | 0.17 | 2.43 | 0.24 | 0.01 |

Parallel model parameter estimates from 3-parameter logistic regression model (Hill). $E_{max}$ represents normalized mean *UBE3A-AS* RNA levels relative to control at 3 μM.
Abbreviations: SEM, standard error of the mean

**Table S4. Chemically Optimized ASOs**

| ASO | RNA Modification | Backbone | Size (nt) | Design (5´-3´) | Sequence (5’-3’) |
|---|---|---|---|---|---|
| 4.0.PO-1.O | OMe | PS/PO | 20 | 5-10-5 | U$^{O}$*A$^{O}$*G$^{O}$*A$^{O}$*A$^{O}$-t*g*g*c*a*c*a*t*c*t-C$^{O}$*U$^{O}$*U$^{O}$*G$^{O}$*G$^{O}$ |
| 4.0.PO-2.O | OMe | PS/PO | 20 | 5-10-5 | U$^{O}$*A$^{O}$-G$^{O}$*A$^{O}$-A$^{O}$*t-g*g-c*a-c*a-t*c-t*C$^{O}$-U$^{O}$*U$^{O}$-G$^{O}$*G$^{O}$ |
| 4.0.PS.M | MOE | PS | 20 | 5-10-5 | T$^{M}$*A$^{M}$*G$^{M}$*A$^{M}$*A$^{M}$*t*g*g*c*a*c*a*t*c*t*5mC$^{M}$*T$^{M}$*T$^{M}$*G$^{M}$*G$^{M}$ |
| 4.0.PO-1.M | MOE | PS/PO | 20 | 5-10-5 | T$^{M}$*A$^{M}$*G$^{M}$*A$^{M}$*A$^{M}$-t*g*g*c*a*c*a*t*c*t-5mC$^{M}$*T$^{M}$*T$^{M}$*G$^{M}$*G$^{M}$ |
| 4.0.PO-2.M | MOE | PS/PO | 20 | 5-10-5 | T$^{M}$*A$^{M}$-G$^{M}$*A$^{M}$-A$^{M}$*t-g*g-c*a-c*a-t*c-t*5mC$^{M}$-T$^{M}$*T$^{M}$-G$^{M}$*G$^{M}$ |
| 4.4.PS.L | LNA | PS | 18 | 3-11-4 | A$^{L}$*G$^{L}$*A$^{L}$*a*t*g*g*c*a*c*a*t*c*t*5mC$^{L}$*T$^{L}$*T$^{L}$*G$^{L}$ |
| 4.4.PO-1.L | LNA | PS/PO | 18 | 3-11-4 | A$^{L}$*G$^{L}$*A$^{L}$-a*t*g*g*c*a*c*a*t*c*t-5mC$^{L}$*T$^{L}$*T$^{L}$*G$^{L}$ |
| 4.4.PO-2.L | LNA | PS/PO | 18 | 3-11-4 | A$^{L}$*G$^{L}$-A$^{L}$*a-t*g-g*c-a*c-a*t-c*t-5mC$^{L}$*T$^{L}$-T$^{L}$*G$^{L}$ |
| 6.1.PO-1.O | OMe | PS/PO | 18 | 4-10-4 | C$^{O}$*U$^{O}$*G$^{O}$*G$^{O}$-t*g*t*c*a*a*c*a*a*g-C$^{O}$*C$^{O}$*A$^{O}$*A$^{O}$ |
| 6.1.PO-2.O | OMe | PS/PO | 18 | 4-10-4 | C$^{O}$*U$^{O}$-G$^{O}$*G$^{O}$-t*g-t*c-a*a-c*a-a*g-C$^{O}$*C$^{O}$-A$^{O}$*A$^{O}$ |
| 6.1.PS.M | MOE | PS | 18 | 4-10-4 | 5mC$^{M}$*T$^{M}$*G$^{M}$*G$^{M}$*t*g*t*c*a*a*c*a*a*g*5mC$^{M}$*5mC$^{M}$*A$^{M}$*A$^{M}$ |
| 6.1.PO-1.M | MOE | PS/PO | 18 | 4-10-4 | 5mC$^{M}$*T$^{M}$*G$^{M}$*G$^{M}$-t*g*t*c*a*a*c*a*a*g-5mC$^{M}$*5mC$^{M}$*A$^{M}$*A$^{M}$ |
| 6.1.PO-2.M | MOE | PS/PO | 18 | 4-10-4 | 5mC$^{M}$*T$^{M}$-G$^{M}$*G$^{M}$-t*g-t*c-a*a-c*a-a*g-5mC$^{M}$*5mC$^{M}$-A$^{M}$*A$^{M}$ |
| 6.2.PS.L | LNA | PS | 17 | 3-10-4 | T$^{L}$*G$^{L}$*G$^{L}$*t*g*t*c*a*a*c*a*a*g*5mC$^{L}$*5mC$^{L}$*A$^{L}$*A$^{L}$ |
| 6.2.PO-1.L | LNA | PS/PO | 17 | 3-10-4 | T$^{L}$*G$^{L}$*G$^{L}$-t*g*t*c*a*a*c*a*a*g-5mC$^{L}$*5mC$^{L}$*A$^{L}$*A$^{L}$ |
| 6.2.PO-2.L | LNA | PS/PO | 17 | 3-10-4 | T$^{L}$*G$^{L}$-G$^{L}$*t-g*t-c*a-a*c-a*a-g*5mC$^{L}$-5mC$^{L}$-A$^{L}$*A$^{L}$ |

Capital letter, RNA; lower-case letter, DNA
Abbreviations: nt, nucleotide; PS and *, phosphorothioate; PO & -, phosphodiester; O, OMe; M, MOE; L, LNA; 5mC, 5-methylcytosine
Note: ASO 4.4 and 6.2 are 18 and 17 nucleotides in length, respectively.

**Table S5. Pharmacological Properties of Chemically Optimized ASOs**

| ASO | $IC_{50}$ (nM) | SEM | Relative Potency | $E_{max}$ (30 µM) | SEM |
|---|---|---|---|---|---|
| 4.0.PS.O | 2,078 | 0.00 | 1.00 | 0.14 | 0.04 |
| 4.0.PO-1.O | 790 | 0.56 | 2.63 | 0.11 | 0.01 |
| 4.0.PO-2.O | 43,588 | 0.01 | 0.05 | 0.61 | 0.03 |
| 4.0.PS.M | 271 | 1.66 | 7.66 | 0.05 | 0.01 |
| 4.0.PO-1.M | 288 | 1.54 | 7.21 | 0.05 | 0.01 |
| 4.0.PO-2.M | 21,031 | 0.02 | 0.10 | 0.50 | 0.03 |
| 4.4.PS.L | 50 | 9.07 | 41.36 | 0.05 | 0.004 |
| 4..4.PO-1.L | 329 | 1.37 | 6.32 | 0.10 | 0.02 |
| 4.4.PO-2.L | 60,311 | 0.01 | 0.03 | 0.61 | 0.03 |
| 6.1.PS.O | 630 | 0.00 | 1.00 | 0.05 | 0.01 |
| 6.1.PO-1.O | 301 | 0.67 | 2.09 | 0.04 | 0.01 |
| 6.1.PO-2.O | 10,030 | 0.02 | 0.06 | 0.25 | 0.03 |
| 6.1.PS.M | 223 | 0.91 | 2.82 | 0.02 | 0.005 |
| 6.1.PO-1.M | 262 | 0.77 | 2.40 | 0.04 | 0.02 |
| 6.1.PO-2.M | 8,696 | 0.02 | 0.07 | 0.31 | 0.03 |
| 6.2.PS.L | 415 | 0.49 | 1.52 | 0.04 | 0.003 |
| 6.2.PO-1.L | 2,050 | 0.10 | 0.31 | 0.17 | 0.03 |
| 6.2.PO-2.L | 7,000 | 0.03 | 0.09 | 0.43 | 0.06 |

Parallel model parameter estimates from 4-parameter logistic regression model (Hill). $E_{max}$ represents normalized mean *UBE3A-AS* RNA levels relative to control at 30 µM.
Abbreviations: SEM, standard error of the mean

| **Table S6. Parental *UBE3A* Allelic Expression in GABAergic Neurons** | | | |
|---|---|---|---|
| **ASO** | **Concentration** | **Paternal *UBEA*** | **Maternal *UBE3A*** |
| **6.1.PO-1.O** | 1 nM | 0.21 | 0.79 |
| | 3 nM | 0.16 | 0.84 |
| | 10 nM | 0.12 | 0.88 |
| | 30 nM | 0.14 | 0.86 |
| | 100 nM | 0.17 | 0.83 |
| | 300 nM | 0.17 | 0.84 |
| | 1 µM | 0.20 | 0.80 |
| | 3 µM | 0.27 | 0.73 |
| | 10 µM | 0.40 | 0.60 |
| | 30 µM | 0.41 | 0.60 |
| **4.4.PS.L** | 1 nM | 0.15 | 0.85 |
| | 3 nM | 0.21 | 0.79 |
| | 10 nM | 0.14 | 0.86 |
| | 30 nM | 0.12 | 0.88 |
| | 100 nM | 0.17 | 0.83 |
| | 300 nM | 0.33 | 0.67 |
| | 1 µM | 0.36 | 0.64 |
| | 3 µM | 0.42 | 0.58 |
| | 10 µM | 0.44 | 0.56 |
| | 30 µM | 0.44 | 0.56 |
| **4.4.PO-1.L** | 1 nM | 0.12 | 0.88 |
| | 3 nM | 0.11 | 0.89 |
| | 10 nM | 0.12 | 0.88 |
| | 30 nM | 0.11 | 0.89 |
| | 100 nM | 0.13 | 0.87 |
| | 300 nM | 0.16 | 0.84 |
| | 1 µM | 0.19 | 0.81 |
| | 3 µM | 0.29 | 0.71 |
| | 10 µM | 0.34 | 0.66 |
| | 30 µM | 0.38 | 0.62 |
| **4.0.PO-1.M** | 1 nM | 0.15 | 0.85 |
| | 3 nM | 0.18 | 0.82 |
| | 10 nM | 0.14 | 0.86 |
| | 30 nM | 0.26 | 0.74 |
| | 100 nM | 0.17 | 0.83 |
| | 300 nM | 0.22 | 0.78 |
| | 1 µM | 0.21 | 0.79 |
| | 3 µM | 0.27 | 0.73 |
| | 10 µM | 0.32 | 0.68 |
| | 30 µM | 0.39 | 0.61 |

**Table S7. Non-Human Primates**

| Animal # | Species | Study | Age (years) | Sex | Treatment | Dose (mg) | Analysis | Analysis 2 |
|---|---|---|---|---|---|---|---|---|
| 1 | *Macaca fascicularis* | 1,2,3 | 3 | M | Vehicle | 0 | *UBE3A-AS* | |
| 2 | *Macaca fascicularis* | 1,2,3 | 3 | F | Vehicle | 0 | *UBE3A-AS* | |
| 3 | *Macaca fascicularis* | 1,2,3 | 2-4 | M | Vehicle | 0 | *UBE3A-AS* | |
| 4 | *Macaca fascicularis* | 1,2,3 | 2-4 | M | Vehicle | 0 | *UBE3A-AS* | |
| 5 | *Macaca fascicularis* | 1,2,3 | 2-4 | M | Vehicle | 0 | *UBE3A-AS* | |
| 6 | *Macaca fascicularis* | 1 | 3 | F | 4.4.PS.L | 10 | *UBE3A-AS* | |
| 7 | *Macaca fascicularis* | 1 | 3 | F | 4.4.PS.L | 10 | *UBE3A-AS* | |
| 8 | *Macaca fascicularis* | 1 | 3 | F | 4.4.PS.L | 10 | *UBE3A-AS* | |
| 9 | *Macaca fascicularis* | 1 | 2-4 | M | 6.1.PO-1.O | 10 | *UBE3A-AS* | |
| 10 | *Macaca fascicularis* | 1 | 2-4 | M | 6.1.PO-1.O | 10 | *UBE3A-AS* | |
| 11 | *Macaca fascicularis* | 1 | 2-4 | M | 6.1.PO-1.O | 10 | *UBE3A-AS* | |
| 12 | *Macaca fascicularis* | 2,3 | 3 | F | Vehicle | 0 | *UBE3A-AS* | |
| 13 | *Macaca fascicularis* | 2 | 3 | F | 4.0.PO-1.M | 10, 10 | *UBE3A-AS* | |
| 14 | *Macaca fascicularis* | 2 | 3 | F | 4.0.PO-1.M | 10, 10 | *UBE3A-AS* | *UBE3A SNP* |
| 15 | *Macaca fascicularis* | 2 | 2-4 | M | 6.1.PO-1.O | 10, 10 | *UBE3A-AS* | *UBE3A SNP* |
| 16 | *Macaca fascicularis* | 2 | 2-4 | M | 6.1.PO-1.O | 10, 10 | *UBE3A-AS* | |
| 17 | *Macaca fascicularis* | 3 | 3 | F | 4.4.PS.L | 7, 3 | *UBE3A-AS* | |
| 18 | *Macaca fascicularis* | 3 | 3 | F | 4.4.PS.L | 7, 3 | *UBE3A-AS* | *UBE3A SNP* |
| 19 | *Macaca fascicularis* | 3 | 3 | F | 4.4.PO-1.L | 7, 3 | *UBE3A-AS* | |
| 20 | *Macaca fascicularis* | 3 | 3 | F | 4.4.PO-1.L | 7, 3 | *UBE3A-AS* | *UBE3A SNP* |
| 21 | *Macaca fascicularis* | 4 | 2-4 | F | Vehicle | 0 | *UBE3A-AS* | |
| 22 | *Macaca fascicularis* | 4 | 2-4 | F | Vehicle | 0 | *UBE3A-AS* | |
| 23 | *Macaca fascicularis* | 4 | 2-4 | F | Vehicle | 0 | *UBE3A-AS* | |
| 24 | *Macaca fascicularis* | 4 | 2-4 | M | 4.4.PS.L | 0.5, 0.5, 0.5 | *UBE3A-AS* | |
| 25 | *Macaca fascicularis* | 4 | 2-4 | F | 4.4.PS.L | 0.5, 0.5, 0.5 | *UBE3A-AS* | |
| 26 | *Macaca fascicularis* | 4 | 2-4 | F | 4.4.PS.L | 0.5, 0.5, 0.5 | *UBE3A-AS* | |
| 27 | *Macaca fascicularis* | 4 | 2-4 | F | 4.4.PS.L | 0.5, 0.5, 0.5 | *UBE3A-AS* | *UBE3A SNP* |
| 28 | *Macaca fascicularis* | 4 | 2-4 | F | 4.4.PS.L | 0.5, 0.5, 0.5 | *UBE3A-AS* | |
| 29 | *Macaca fascicularis* | 4 | 2-4 | M | 4.4.PS.L | 3, 3, 3 | *UBE3A-AS* | |
| 30 | *Macaca fascicularis* | 4 | 2-4 | M | 4.4.PS.L | 3, 3, 3 | *UBE3A-AS* | |
| 31 | *Macaca fascicularis* | 4 | 2-4 | F | 4.4.PS.L | 3, 3, 3 | *UBE3A-AS* | *UBE3A SNP* |
| 32 | *Macaca fascicularis* | 4 | 2-4 | F | 4.4.PS.L | 3, 3, 3 | *UBE3A-AS* | |
| 33 | *Macaca fascicularis* | 4 | 2-4 | F | 4.4.PS.L | 3, 3, 3 | *UBE3A-AS* | |
| 34 | *Macaca fascicularis* | 4 | 2-4 | M | 4.4.PS.L | 5, 5, 5 | *UBE3A-AS* | *UBE3A SNP* |
| 35 | *Macaca fascicularis* | 4 | 2-4 | M | 4.4.PS.L | 5, 5, 5 | *UBE3A-AS* | |
| 36 | *Macaca fascicularis* | 4 | 2-4 | F | 4.4.PS.L | 5, 5, 5 | *UBE3A-AS* | *UBE3A SNP* |
| 37 | *Macaca fascicularis* | 4 | 2-4 | F | 4.4.PS.L | 5, 5, 5 | *UBE3A-AS* | |
| 38 | *Macaca fascicularis* | 4 | 2-4 | F | 4.4.PS.L | 5, 5, 5 | *UBE3A-AS* | |
| 39 | *Macaca fascicularis* | 4 | 2-4 | F | 4.4.PO-1.L | 0.5, 0.5, 0.5 | *UBE3A-AS* | |
| 40 | *Macaca fascicularis* | 4 | 2-4 | F | 4.4.PO-1.L | 0.5, 0.5, 0.5 | *UBE3A-AS* | |
| 41 | *Macaca fascicularis* | 4 | 2-4 | F | 4.4.PO-1.L | 0.5, 0.5, 0.5 | *UBE3A-AS* | |
| 42 | *Macaca fascicularis* | 4 | 2-4 | F | 4.4.PO-1.L | 3, 3, 3 | *UBE3A-AS* | |
| 43 | *Macaca fascicularis* | 4 | 2-4 | F | 4.4.PO-1.L | 3, 3, 3 | *UBE3A-AS* | |
| 44 | *Macaca fascicularis* | 4 | 2-4 | F | 4.4.PO-1.L | 3, 3, 3 | *UBE3A-AS* | |
| 45 | *Macaca fascicularis* | 4 | 2-4 | F | 4.4.PO-1.L | 5, 5, 5 | *UBE3A-AS* | |
| 46 | *Macaca fascicularis* | 4 | 2-4 | F | 4.4.PO-1.L | 5, 5, 5 | *UBE3A-AS* | |
| 47 | *Macaca fascicularis* | 4 | 2-4 | F | 4.4.PO-1.L | 5, 5, 5 | *UBE3A-AS* | *UBE3A SNP* |
| 48 | *Macaca fascicularis* | NA | 3 | M | Vehicle | 0 | NA | *UBE3A SNP* |

**Table S8. Parental *UBE3A* Allelic Expression by Tissue**

| Tissue | Animal # | Treatment | Dose (mg) | Total Dose (mg) | Paternal *UBE3A* | Maternal *UBE3A* |
|---|---|---|---|---|---|---|
| **Motor Cortex** | 48 | Vehicle | 0 | 0 | 11.40 | 88.60 |
| | 14 | 4.0.PO-1.M | 10,10 | 20 | 19.10 | 80.90 |
| | 15 | 6.1.PO-1.O | 10,10 | 20 | 16.85 | 83.15 |
| | 20 | 4.4.PO-1.L | 7,3 | 10 | 38.30 | 61.70 |
| | 47 | 4.4.PO-1.L | 5, 5, 5 | 15 | 21.05 | 78.95 |
| | 27 | 4.4.PS.L | 0.5, 0.5, 0.5 | 1.5 | 28.30 | 71.70 |
| | 31 | 4.4.PS.L | 3, 3, 3 | 9 | 23.55 | 76.45 |
| | 18 | 4.4.PS.L | 7,3 | 10 | 24.65 | 75.35 |
| | 34 | 4.4.PS.L | 5, 5, 5 | 15 | 38.00 | 62.00 |
| | 36 | 4.4.PS.L | 5, 5, 5 | 15 | 34.90 | 65.10 |
| **Frontal Cortex** | 48 | Vehicle | 0 | 0 | 13.12 | 86.88 |
| | 14 | 4.0.PO-1.M | 10,10 | 20 | 17.60 | 82.40 |
| | 15 | 6.1.PO-1.O | 10,10 | 20 | 18.30 | 81.70 |
| | 20 | 4.4.PO-1.L | 7,3 | 10 | 31.60 | 68.40 |
| | 47 | 4.4.PO-1.L | 5, 5, 5 | 15 | 20.80 | 79.20 |
| | 27 | 4.4.PS.L | 0.5, 0.5, 0.5 | 1.5 | 26.60 | 73.40 |
| | 31 | 4.4.PS.L | 3, 3, 3 | 9 | 32.60 | 67.40 |
| | 18 | 4.4.PS.L | 7,3 | 10 | 33.10 | 66.90 |
| | 34 | 4.4.PS.L | 5, 5, 5 | 15 | 35.10 | 64.90 |
| | 36 | 4.4.PS.L | 5, 5, 5 | 15 | 39.40 | 60.60 |
| **Hippocampus** | 48 | Vehicle | 0 | 0 | 7.68 | 92.32 |
| | 14 | 4.0.PO-1.M | 10,10 | 20 | 9.10 | 90.90 |
| | 15 | 6.1.PO-1.O | 10,10 | 20 | 12.90 | 87.10 |
| | 20 | 4.4.PO-1.L | 7,3 | 10 | 9.13 | 90.87 |
| | 47 | 4.4.PO-1.L | 5, 5, 5 | 15 | 16.10 | 83.90 |
| | 27 | 4.4.PS.L | 0.5, 0.5, 0.5 | 1.5 | 15.65 | 84.35 |
| | 31 | 4.4.PS.L | 3, 3, 3 | 9 | 15.30 | 84.70 |
| | 18 | 4.4.PS.L | 7,3 | 10 | 25.80 | 74.20 |
| | 34 | 4.4.PS.L | 5, 5, 5 | 15 | 22.40 | 77.60 |
| | 36 | 4.4.PS.L | 5, 5, 5 | 15 | 21.35 | 78.65 |
| **LSC** | 48 | Vehicle | 0 | 0 | 30.20 | 69.80 |
| | 14 | 4.0.PO-1.M | 10,10 | 20 | 30.90 | 69.10 |
| | 15 | 6.1.PO-1.O | 10,10 | 20 | 48.50 | 51.50 |
| | 20 | 4.4.PO-1.L | 7,3 | 10 | 36.20 | 63.80 |
| | 47 | 4.4.PO-1.L | 5, 5, 5 | 15 | 46.90 | 53.10 |
| | 27 | 4.4.PS.L | 0.5, 0.5, 0.5 | 1.5 | 49.70 | 50.30 |
| | 31 | 4.4.PS.L | 3, 3, 3 | 9 | 50.90 | 49.10 |
| | 18 | 4.4.PS.L | 7,3 | 10 | 39.00 | 61.00 |
| | 34 | 4.4.PS.L | 5, 5, 5 | 15 | 50.30 | 49.70 |
| | 36 | 4.4.PS.L | 5, 5, 5 | 15 | 50.40 | 49.60 |

Data represent expression (fractional abundance) of paternal and maternal *UBE3A* RNA levels.
Abbreviations: LSC, lumbar spinal cord

#### Table S9. Parental *UBE3A* Allelic Expression by Animal

| Animal # | Treatment | Dose (mg) | Total Dose (mg) | Paternal *UBE3A* | Maternal *UBE3A* |
|---|---|---|---|---|---|
| 48 | Vehicle | 0 | 0 | 15.60 | 84.40 |
| 14 | 4.0.PO-1.M | 10,10 | 20 | 19.16 | 80.84 |
| 15 | 6.1.PO-1.O | 10,10 | 20 | 22.68 | 77.32 |
| 20 | 4.4.PO-1.L | 7,3 | 10 | 30.71 | 69.29 |
| 47 | 4.4.PO-1.L | 5, 5, 5 | 15 | 25.18 | 74.82 |
| 27 | 4.4.PS.L | 0.5, 0.5, 0.5 | 1.5 | 27.37 | 72.63 |
| 31 | 4.4.PS.L | 3, 3, 3 | 9 | 26.87 | 73.13 |
| 18 | 4.4.PS.L | 7,3 | 10 | 29.44 | 70.56 |
| 34 | 4.4.PS.L | 5, 5, 5 | 15 | 34.37 | 65.63 |
| 36 | 4.4.PS.L | 5, 5, 5 | 15 | 33.72 | 66.28 |

Data represent the mean *UBE3A* allelic expression (fractional abundance) across the motor cortex, frontal cortex, hippocampus, and lumbar spinal cord.

| Table S10. Key Resources | | |
|---|---|---|
| REAGENT or RESOURCE | SOURCE | IDENTIFIER |
| **Antibodies** | | |
| Purified Mouse Anti-UBE3A | BD BioSciences | Cat# BD611416; RRID:AB_2338692 |
| HRP - Goat Anti-mouse IgG (H+L) | ThermoFisher Scientific | Cat# A16066; RRID:AB_2534739 |
| Qiagen RNAeasy Plus Kit | Qiagen | Cat# 74136 |
| Gene Expression Master Mix | ThermoFisher Scientific | Cat# 4369016 |
| Illumina TruSeq Stranded Total RNA Kit | Illumina | Cat# 20020597 |
| Illumina TruSeq Stranded mRNA Kit | Illumina | Cat# 20020595 |
| D1000 DNA ScreenTape | Agilent Technologies | Cat# 5067-5582 |
| D1000 Reagents | Agilent Technologies | Cat# 5067-5583 |
| RNA ScreenTape | Agilent Technologies | Cat# 5067-5576 |
| RNA ScreenTape Sample Buffer | Agilent Technologies | Cat# 5067-5577 |
| Laminin | ThermoFisher Scientific | Cat# 23017-015 |
| Neurobasal A Medium | ThermoFisher Scientific | Cat# 10888022 |
| B27 | ThermoFisher Scientific | Cat# 17504044 |
| Topotecan | Millipore Sigma | Cat# T2705 |
| 20% Knockout Serum Replacement | ThermoFisher Scientific | Cat# 10828-028 |
| PluriSTEM Human ES/iPS Medium | Millipore Sigma | Cat# SCM130 |
| Dispase II | Millipore Sigma | Cat# SCM133 |
| Matrigel hESC-qualified Matrix | Corning | Cat# 354277 |
| Vitronectin | Millipore Sigma | Cat# CC130 |
| Millipore ES/iPS Neurogenesis Kit | Millipore Sigma | Cat# SCR603, SCM110, SCM111 |
| EZ-LiFT | Millipore Sigma | Cat# SCM139 |
| ENStem-A Neural Expansion Medium | Millipore Sigma | Cat# SCM004 |
| Differentiation Medium | Millipore Sigma | Cat# SCM111 |
| Cellular Dynamics Maintenance Medium | Cellular Dynamics | Cat# NRM-100-121-001 |
| Cells-to-CT kit | ThermoFisher Scientific | Cat# AM1728 |
| 4x Laemmli Buffer | Bio-Rad | Cat# 1610747 |
| 7.5% Mini Protean TGX Stain-Free Protein Gels | Bio-Rad | Cat# 4568024 |
| 10X TGS Buffer | Bio-Rad | Cat# 1610772 |
| Clarity Western ECL Substrates | Bio-Rad | Cat# 170-5061 |
| ddPCR Supermix for Probes (no dUTP) | Bio-Rad | Cat# 1863024 |
| Qubit RNA BR Assay Kit | ThermoFisher Scientific | Cat# Q10211 |
| Qubit DNA HS Assay Kit | ThermoFisher Scientific | Cat# Q32851 |
| High-Capacity RNA-to-cDNA kit | ThermoFisher Scientific | Cat# 4387406 |
| Dneasy Blood and Tissue Kit | Qiagen | Cat# 69504 |
| 2x Phire Hot Start II PCR master mix | ThermoFisher Scientific | Cat# F126L |
| PureLink Quick Gel Extraction & PCR Purification Combo Kit | ThermoFisher Scientific | Cat# K220001 |
| PowerUp SYBR Green Master Mix | ThermoFisher Scientific | Cat# A25779 |
| Trans-Blot Turbo Mini NitroCellulose Transfer Packs | Bio-Rad | Cat# 1704158 |
| DMEM/F12 | ThermoFisher Scientific | Cat# 11330-057 |
| SuperScript IV First-Strand Synthesis System | ThermoFisher Scientific | Cat# 18091050 |
| Growth Factor Reduced Matrigel Matrix | Corning | Cat# 354230 |
| Human Cortex | Agilent Technologies | Cat# 540005 |
| aCSF | Harvard Apparatus | Cat# 59-7316 |
| Human PPIA | ThermoFisher Scientific | Cat# Hs99999904_m1 |
| Human UBE3A-AS | ThermoFisher Scientific | Cat# Hs01372957_m1 |
| Human SNORD116-11 | ThermoFisher Scientific | Cat# Hs04275268_gH |
| Human SNORD115 | ThermoFisher Scientific | Cat# Hs04275288_gH |
| Human IPW/116HG | ThermoFisher Scientific | Cat# Hs03455409_s1 |
| Macaque PPIA | ThermoFisher Scientific | Cat# Mf04932064_gH |
| GABA iPSC neural precursor cells | Cellular Dynamics | Cat# NRC-100-010-001 |
| AG1-0 iPSCs | Kerafast | Cat# ECN001 |
| Female and Male Cynomolgus Monkeys | Charles River Laboratories and Northern Biomedical Research Inc | |
| C57BL/6J Mice | The Jackson Laboratory | Cat# JAX:000664, RRID:IMSR_JAX:000664 |
| RepeatMasker | | (http://www.repeatmasker.org) |

| | | |
|---|---|---|
| Soligo | Software for Statistical Folding of Nucleic Acid and Studies of Regulatory RNA | http://sfold.wadsworth.org |
| Image J | NIH | |
| Quantasoft Software | Bio-Rad | |
| Image Lab 5.2.1 | Bio-Rad | |
| BioRad CFX Maestro Software | Bio-Rad | |
| JMP Pro 15 | | SAS Institute Inc. |
| RNAfold | | http://rna.tbi.univie.ac.at/cgi-bin/RNAWebSuite/RNAfold.cgi |
| Mfold | | http://unafold.rna.albany.edu/?q=mfold/RNA-Folding-Form |
| EMBOSS | | https://www.ebi.ac.uk/Tools/psa/ |
| CLC Genomics Workbench | Qiagen | v8.0.1 |
| UCSC Genome Browser | | http://genome.ucsc.edu/ |
| RepeatMasker | Smit AFA, Hubley R, Green P. RepeatMasker Open-3.0 | http://www.repeatmasker.org |
| RNA Sequencing Data | Agilent Technologies | Cat# SLGV013SL |
| NextSeq 500 | Illumina Inc. | https://genomics.tamu.edu/genomics-core/ |
| NovaSeq 6000 | Illumina Inc. | https://northtexasgenomecenter.com |
| bcl2fastq v2.20.0.422 | Illumina Inc. | https://support.illumina.com/sequencing/sequencing_software/bcl2fastq-conversion-software.html |
| 0.2-µm Millex GV PVDF syringe filter | Millipore Sigma | Cat# SLGV013SL |
| MINLOVOL | Instech Solomon | |
